# Dopamine biases sensitivity to personal goals and social influence in self-control over everyday desires

**DOI:** 10.1101/2021.09.10.459829

**Authors:** Jaime J. Castrellon, Jacob S. Young, Linh C. Dang, Christopher T. Smith, Ronald L. Cowan, David H. Zald, Gregory R. Samanez-Larkin

## Abstract

People regularly give in to daily temptations in spite of conflict with personal goals. To test hypotheses about neuropharmacological influences on self-control, we used positron emission tomography to measure dopamine D2-like receptors (D2R) and experience sampling surveys to naturalistically track daily desires outside the laboratory in everyday life in a sample of 103 adults. Higher D2R availability in the ventral striatum was associated with increased sensitivity to personal goal conflict but not desire strength in deciding whether to attempt to resist a desire. The influence of D2Rs on sensitivity to personal goal conflict depended on whether desires were experienced in a social context. D2R availability in the midbrain (but not the ventral striatum) influenced whether desires were enacted. These findings provide unique evidence that the dopamine system influences decision making and regulatory behavior and provides new insights into how these mechanisms interact with personal goals and social contexts.

## Introduction

To successfully navigate everyday life, humans regularly set long-term personal goals like performing well at work or school or losing weight. In pursuit of these goals, humans also experience and naturally enact conflicting desires like watching Netflix and scrolling Twitter or consuming sweets. Integrating the benefits of personal goals with immediate desires is central to self-control (Fujita, 2011). What, then, makes some people more sensitive to their personal goals or immediate desires? The neurotransmitter dopamine (DA) has been speculated to impact individual differences in self-control abilities. Specifically, DA D2-like receptors (D2Rs) shape the ability to inhibit actions in well-controlled tasks in the laboratory (Frank, 2005; Ghahremani et al., 2012; Robertson et al., 2015). However, no study to date has examined whether variation in D2Rs biases decisions about, or the successful execution of, self-control as desires are spontaneously experienced in daily life.

Self-control in the context of regulating desires in everyday life has been conceptualized as a process that involves generating control motivation based on the strength of a desire and level of personal goal conflict. Control motivation is the “aspiration to control desire” and shapes how much effort a person allocates in the decision process (Kotabe & Hofmann, 2015). The result of control motivation and effort allocation in desire resistance can manifest as a conscious resistance attempt. This kind of mental effort motivation in support of resistance attempts, however, may also depend on additional cues in the environment (Shenhav et al., 2021). Cues from the behaviors of others, for example, have been long-known to influence conformity in decision making (Asch, 1955; Cialdini & Goldstein, 2004). While susceptibility to peer conformity pressures are often highlighted in adolescence, they diminish but do not disappear in adulthood (Knoll et al., 2015; Steinberg & Monahan, 2007). Work using the same experimental paradigm that we use here has also shown that the presence of others engaging in a desire lowers attempts to resist those desires (Hofmann, Baumeister, et al., 2012). In that experiment, the authors attributed social influence effects on desire resistance to the possibility that participants engage in self-justification and motivated reasoning, leading them to feel more comfortable in co-enacting a desire (Kivetz & Zheng, 2006) and behavioral mimicry facilitating motor systems to mirror actions of others (Chartrand & Bargh, 1999). It is possible that the importance of personal goals (not just attempts to resist desires) could also depend on the presence of others enacting a desire. Observing others enacting a desire may momentarily attenuate the value of long-term goals if doing so lowers the risk of social rejection and supports building and maintaining relationships (Cullum et al., 2011). This view is consistent with work showing that reward valuation is malleable to social influence (Bixter & Rogers, 2019; Zaki et al., 2011). This malleability of the importance of conflicting personal goals to self-regulation could be adaptive for social well-being at the cost of lowering resistance to temptations.

Laboratory studies suggest that features of the DA system could shape decisions to practice self-control over everyday desires. In addition to its influence on the ability to inhibit responses (Ghahremani et al., 2012; Robertson et al., 2015), DA has been shown to impact healthy adults’ decisions about willingness to expend effort (Treadway et al., 2012), and the underlying neural representations of the subjective value of benefits at the cost of delays and effort (Castrellon et al., 2019; Medic et al., 2014; Pine et al., 2010). D2Rs may similarly impact attempts to resist desires by biasing the potential benefits of personal goals over immediate temptations, and the willingness to exert self-control effort. Differences in several self-control-related personality traits have been linked to D2R availability in regions such as the ventral striatum (Caravaggio et al., 2016; Reeves et al., 2012), amygdala (Okita et al., 2016), and midbrain (Buckholtz et al., 2010; Zald et al., 2008). However, no study to date has directly measured the dopamine system in humans and evaluated associations with self-control in everyday life.

While research in humans has not yet identified whether DA signaling in specific regions influences sensitivity to social contexts, pharmacological manipulation studies indicate that DA biases the calculation of social costs related to harm (Crockett et al., 2015) and prosocial giving (Sáez et al., 2015) and increases sensitivity to subtle social emotional expressions (Wardle et al., 2012). Further, higher social attachment has been shown to be negatively correlated with D2R availability in humans (Caravaggio et al., 2017), indicating a potential role for DA in mediating sensitivity to social information. Additional work in other animals corroborates these associations showing that D2Rs affect prairie vole pair bonding (Wang et al., 1999) and that dopamine neurons in rats encode prediction errors that support social learning from conspecifics (Prévost-Solié et al., 2020). Together, these findings suggest that DA could differentially affect sensitivity to the perceived benefits and costs of self-control when people are in the presence of others enacting a desire.

The present study sought to identify whether individual differences in D2R availability are related to differences in how people weight personal goals with immediate temptations in self-control over everyday desires and whether these associations are sensitive to social contexts. Since people experience a diversity of desires multiple times a day that vary in strength and often conflict with a range of personal goal types, we used an experience sampling approach that provided a more ecologically valid assessment of variation in real-world self-control decisions than can be accomplished in an experimental laboratory procedure.

## Methods

103 healthy adults across the adult life span (ages 18-80, M = 35.9, SD = 11.7, 53 women) across 3 study samples underwent a positron emission tomography (PET) scan with the radiotracer [18F]fallypride, which labels DA D2Rs (See Supplemental materials for a description of PET methodology and **Table S1** for participant demographics). These participants also received surveys delivered via text message three times per day for 10 days as part of an experience sampling protocol adapted from (Hofmann, Baumeister, et al., 2012; Hofmann, Vohs, et al., 2012). In each survey, participants indicated up to three desires they had experienced in the past three hours along with: (1) the strength of the desire (0: No desire at all to 7: Irresistible), (2) the degree to which the desire conflicted with a personal goal (0: No conflict at all to 4: Very high conflict), (3) whether they attempted to resist the desire (Yes/No) and (4) whether they enacted those desires (Yes/No). Participants also identified whether others present (either physically or over media) were engaging in the desire (Yes/No). *See Supplemental Material for further details on experience sampling protocol*.

**Fig. 1.**
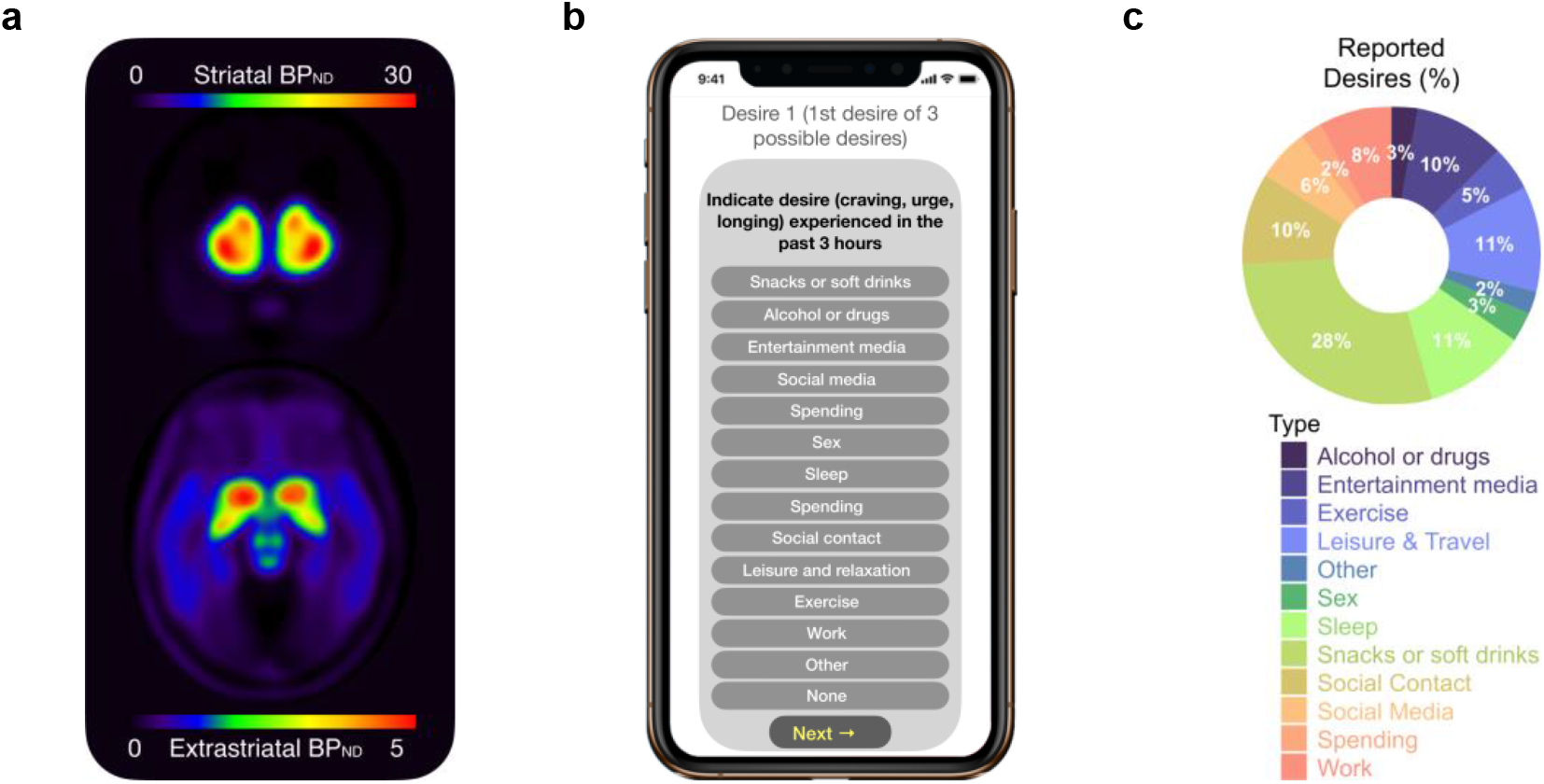
Study methods and reported desires. (**a)** Participants underwent a PET scan with the radiotracer [18F]fallypride to label D2 receptors (D2Rs). The image displays mean D2R availability in the striatum (top, coronal slice) as well as the amygdala and midbrain (bottom, axial slice). (**b)** These same participants completed an experience sampling study in which they completed mobile surveys to indicate desires experienced over the course of 10 days along with the degree to which the desire conflicted with personal goals, whether they attempted to resist and enact those desires, and whether others were present enacting the desires. **(c)** Across participants, the top five most frequently experienced desires included snacking (28%), sleep (11%), leisure and travel (11%), entertainment (10%), and social contact (10%).

To test predictors of self-control, we used multi-level logistic regression with random intercepts for participants (to account for inter-individual variability). The dependent variable was resistance attempt (attempt versus no attempt) and, separately, resistance success (note we use the term resistance success for all cases in which the person did not enact the desire, regardless of whether the person reported a conscious effort to resist the desire). We report standardized regression coefficients (log odds) of predictor variables of interest and use these measures to capture sensitivity to goals and desires as their slope for binary outcome measures. All analyses controlled for participant age, sex, study sample, and the number of days between the PET scan and experience sampling start date. *See Supplemental Material for detailed model information and statistical analysis*.

Prior to advanced modelling, we conducted simple correlation analyses between D2R availability and participants summarized untransformed ratings and decisions. The results of these correlations are reported in **Table S2**. To limit the number of statistical tests for associations with DA, we performed an initial model comparison to evaluate how participants integrate personal goal conflict and desires in self-control decisions. Specifically, we compared models that assume ratings are integrated: (1) linearly as the difference between goal conflict and desire ratings, (2) as a ratio of goal conflict/desire ratings, or (3) independently as distinct weights. Measures of model fit (**Table S3**) indicated that a model based on independent contributions of goal conflict and desire ratings best explained decisions to attempt to resist desires. This model indicated that sensitivity to personal goal conflict increases the odds of attempting to resist a desire and that, separately, desire strength reduces the odds of attempting to resist a desire. We used this model for both resistance attempts and resistance success. Follow-up models evaluated whether social context (being around others enacting a desire) impacts sensitivity of personal goal conflict and desire strength on resistance attempts and success.

To investigate the role of D2R availability on resistance attempts, we tested the independent interactions between either personal goal conflict or desire strength with average D2R availability from three regions of interest (ROI): the ventral striatum, midbrain, and amygdala. These regions were selected because D2R availability in these regions have been previously linked to self-control related behaviors and traits in humans (Buckholtz et al., 2010; Caravaggio et al., 2016; Okita et al., 2016; Reeves et al., 2012; Zald et al., 2008). Resistance success was tested via the interaction between either personal goal conflict or desire strength with DA D2R availability and resistance attempt. To facilitate interpretation of statistically significant cross-level interactions, we conducted simple slopes analysis to identify estimates for participants with low (−1 SD), average, and high (+1 SD) D2R availability. Follow-up analyses tested the effect of social context on self-control with models that included interactions between D2R availability, personal goal conflict or desire strength ratings, and whether others present were also enacting in the desire.

On average, across the 10 days of experience sampling, participants reported 55.86 (SD = 20.19) individual desires, of which they attempted to resist 18.9 desires (SD = 13.59). The average desire strength rating was 4.26 (SD = 0.27) out of 7 and the average personal goal conflict was rated 0.95 (SD = 0.40) out of 4. The entire dataset included 5,752 observations of experienced desires for analyses. (*See Supplemental Material for details on model specification, participant descriptive statistics, and summary-level associations between DA D2R availability and experience sampling data*).

## Results

Behaviorally, attempts to resist desires were strongly predicted by personal goal conflict (β = 1.29, CI [1.21, 1.37], Z = 31.9, p < 0.001) and negatively predicted by desire strength (β = - 0.31, CI [−0.38, −0.23], Z = −8.24, p < 0.001). Full model results including comparison with other models are presented in **Table S3**. Resistance success was predicted by personal goal conflict (β = 0.209, CI [0.06, 0.360], Z = 2.80, p = 0.005), negatively by desire strength (β = −0.322, CI [-0.440, −0.207], Z = −5.50, p < 0.001), resistance attempt (β = 3.26, CI [3.05, 3.48], Z = 30.20, p < 0.001), and the interaction between resistance attempt and personal goal conflict (β = 0.235, CI [0.054, 0.416], Z = 2.55, p = 0.011) but not the interaction between resistance attempt and desire strength (β = −0.074, CI [−0.235, 0.087], Z = −0.902, p = 0.367). Full model results including comparison with other models are shown in **Table S4**.

To examine the impact of social context (the presence of others enacting a desire), we first modeled the two-way interactions between social context and personal goal conflict and social context and desire strength. In addition to main effects of personal goal conflict (β = 1.26, CI [1.15, 1.36], Z = 24.0, p < 0.001) and desire strength (β = −0.27, CI [−0.36, −0.18], Z = −5.70, p < 0.001), this model identified a significant main effect of social context (β = −1.00, CI [−1.16, − 0.18], Z = 2-12.1, p < 0.001). The presence of others enacting a desire lowered the odds of attempting to resist a desire. However, neither the interaction between personal goal conflict and social context (β = −0.01, CI [−0.17, 0.14], Z = −0.15, p = 0.884) nor the interaction between desire strength and social context (β = −0.08, CI [−0.24, 0.07], Z = −1.07, p = 0.287) were statistically significant, indicating that social context did not substantially influence how participants weigh benefits and costs (full model results are shown in **Table S5**). For resistance success, we modeled the three-way interaction between resistance attempt, desire rating (personal goal conflict or desire strength), and social context. Here there was a main effect of social context (β = −1.27, CI [−1.56, −0.99], Z = −8.71, p < 0.001). Although we observed the same two-way interactions between resistance attempt and ratings reported above, there were no other statistically significant two-way or three-way interactions with social context (full model results are shown in **Table S6**). These effects largely replicate prior reports on the impact of personal goal conflict, desires, and social context on self-control (Hofmann, Baumeister, et al., 2012).

Dopamine D2R receptors were unrelated to the number of desires reported or desire strength (See **Table S2**). However, there was a main effect of ventral striatal D2R availability on attempts to resist desires. Higher D2R availability in the ventral striatum was associated with increased resistance attempts (β = 0.296, CI [0.024, 0.567], Z = 2.14, p = 0.032) (**Fig. 2a**). Specifically, with each 1-SD unit increase in receptor availability, the odds of attempting to resist a desire increased by 0.34 (exp(0.296) = 1.34). There were no significant main effects of D2R availability in the midbrain (β = 0.07, CI [−0.25, 0.39], Z = 0.433, p = 0.665) or amygdala (β = 0.210, CI [−0.08, 0.499], Z = 1.42, p = 0.155) on attempts to resist desires. Resistance attempts were predicted by the interaction between personal goal conflict and D2R availability in the ventral striatum (β = 0.15, CI [0.07, 0.23], Z = 3.71, p < 0.001) (**Fig. 2b**), but not the midbrain (β = 0.02, CI [−0.06, 0.10], Z = 0.45, p = 0.656), or amygdala (β = 0.04, CI [−0.05, 0.12], Z = 0.85, p = 0.397). Simple slopes analysis indicated that individuals with high D2R availability (+1 SD) in the ventral striatum were more sensitive to personal goal conflict (β = 1.46, CI [1.33, 1.58], Z = 23.5, p < 0.001) than those with average (β = 1.30, CI [1.22, 1.39], Z = 31.8, p < 0.001) or low D2R availability (−1 SD: β = 1.15, CI [1.05, 1.26], Z = 21.7, p < 0.001). In other words, whereas individuals with low D2R availability were ~2.15 times more likely to attempt to resist a desire with each 1-SD unit increase in personal goal conflict (exp(1.15) = 3.16), those with high D2R availability were ~3.3 times more likely to attempt to resist a desire with each 1-SD unit increase in personal goal conflict (exp(1.46) = 4.31). Resistance attempts were not predicted by the interaction between desire strength and D2R availability in the ventral striatum (β = −0.06, CI [−0.13, 0.01], Z = −1.66, p = 0.096) (**Fig. 2c**), midbrain (β = −0.016, CI [−0.09, 0.06], Z = −0.42, p = 0.674), or amygdala (β = −0.05, CI [−0.12, 0.03], Z = −1.29, p = 0.197). Full model results for interactions effects of D2R availability (in each ROI) on resistance attempts are shown in **Table S7**.

**Fig. 2.**
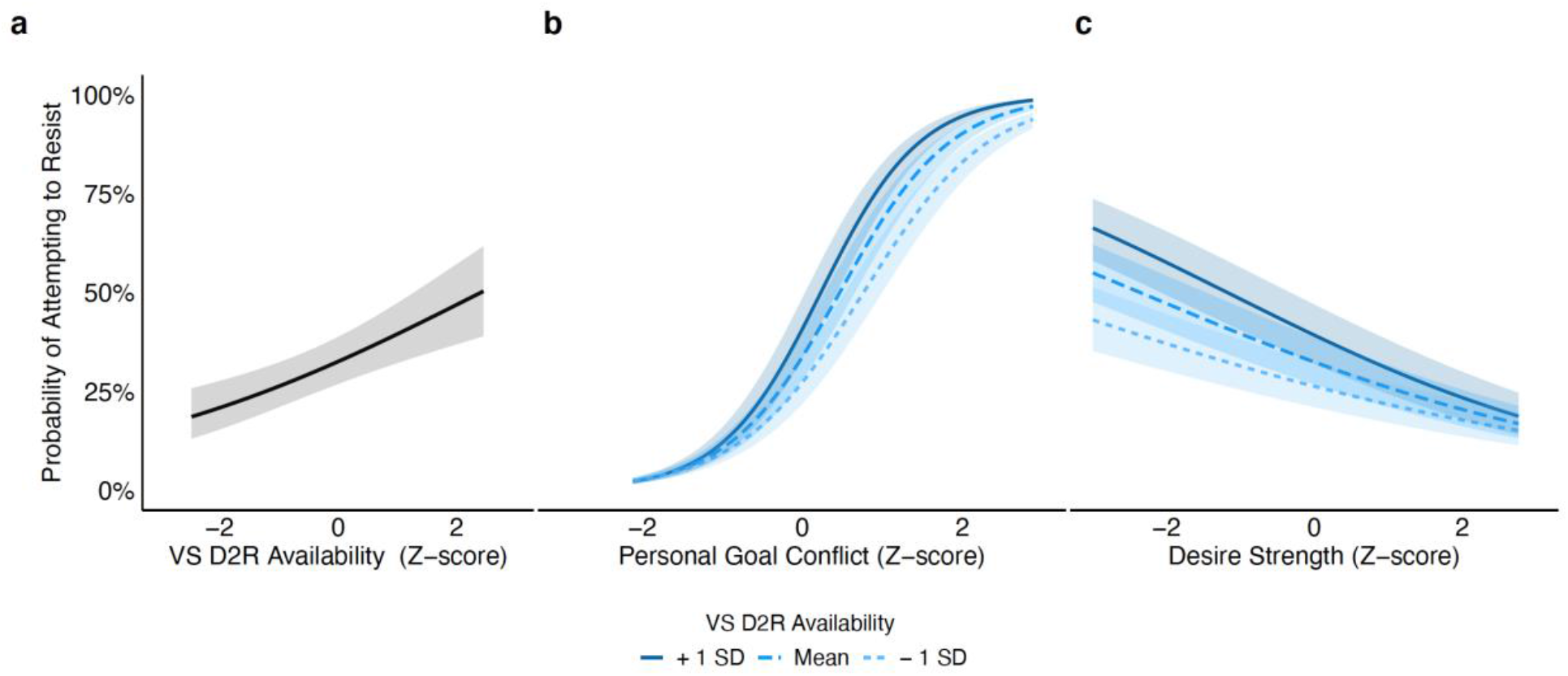
Relationship between dopamine D2 receptor availability and attempts to resist spontaneously experienced desires. Across participants, **(a)** higher D2R availability predicted a higher probability of desire resistance attempt. Personal goal conflict and desire strength ratings are standardized within participants’ survey data and VS D2R availability is standardized across participants. Participants with higher dopamine D2 receptor (D2R) availability in the ventral striatum (VS) were **(b)** more sensitive to conflicting personal goals such that participants with higher D2R availability were more likely to attempt to resist desires as personal goal conflict increased. However, variation in dopamine D2R availability was **(c)** not related to sensitivity to the strength of their desires. Shaded areas represent ± 1 standard error.

To test the hypothesis that D2R influence on resistance attempts is sensitive to social context, we considered whether the presence of others engaging in the desire shifts decisions to attempt to resist desires. Specifically, we tested the three-way interaction between D2R availability, degree of personal goal conflict or desire strength, and whether others were present enacting the desire. For the ventral striatum, in addition to the significant main effects of personal goal conflict, desire strength, and D2R availability, there was a significant main effect of social context (β = −1.00, CI [−1.16, −0.84], Z = −12.01, p < 0.001) indicating that on average, all participants were less likely to attempt to resist a desire when in the presence of others enacting the desire. There were no statistically significant two-way interactions between (1) social context and personal goal conflict (β = −0.04, CI [−0.20, 0.11], Z = −0.55, p = 0.58), (2) social context and desire strength (β = −0.08, CI [−0.23, 0.08], Z = −0.98, p = 0.33), or (3) social context and D2R availability (β = −0.06, CI [−0.21, 0.10], Z = −0.73, p = 0.47). However, there was a statistically significant three-way interaction between social context, personal goal conflict, and VS D2R (β = −0.20, CI [−0.36, −0.04], Z = −2.45, p = 0.014) (**Fig 3**). Simple slopes analysis indicated that participants with high D2R availability were substantially more sensitive to their personal goal conflict (β = 1.52, CI [1.35, 1.68], Z = 18.09, p < 0.001) than those with average (β = 1.29, CI [1.18, 1.39], Z = 23.85, p < 0.001) or low (β = 1.06, CI [0.925, 1.19], Z = 15.8, p < 0.001) D2R availability when not in the presence of others enacting a desire. These differential sensitivities to personal goal conflict changed when desires were experienced in the presence of others engaging in the desire. Specifically, in the presence of others, participants with high D2R availability no longer showed an enhanced sensitivity to their personal goal conflict (β = 1.28, CI [1.10, 1.45], Z = 14.0, p < 0.001) compared to those with average (β = 1.24, CI [1.12, 1.37], Z = 19.9, p < 0.001) or low (β = 1.21, CI [1.04, 1.38], Z = 14.0, p < 0.001) D2R availability. Expressed differently, participants with low D2R availability became more sensitive to their personal goal conflict in a social context in which others were present enacting the desire and that conversely, participants with high D2R availability became less sensitive to their personal goal conflict when in the presence of others enacting the desire. No two-way or three-way interactions with social context were statistically significant for D2R availability in the amygdala or midbrain. Full model results for interactions effects of D2R availability (in each ROI) on resistance attempts are shown in **Table S9**.

**Fig. 3.**
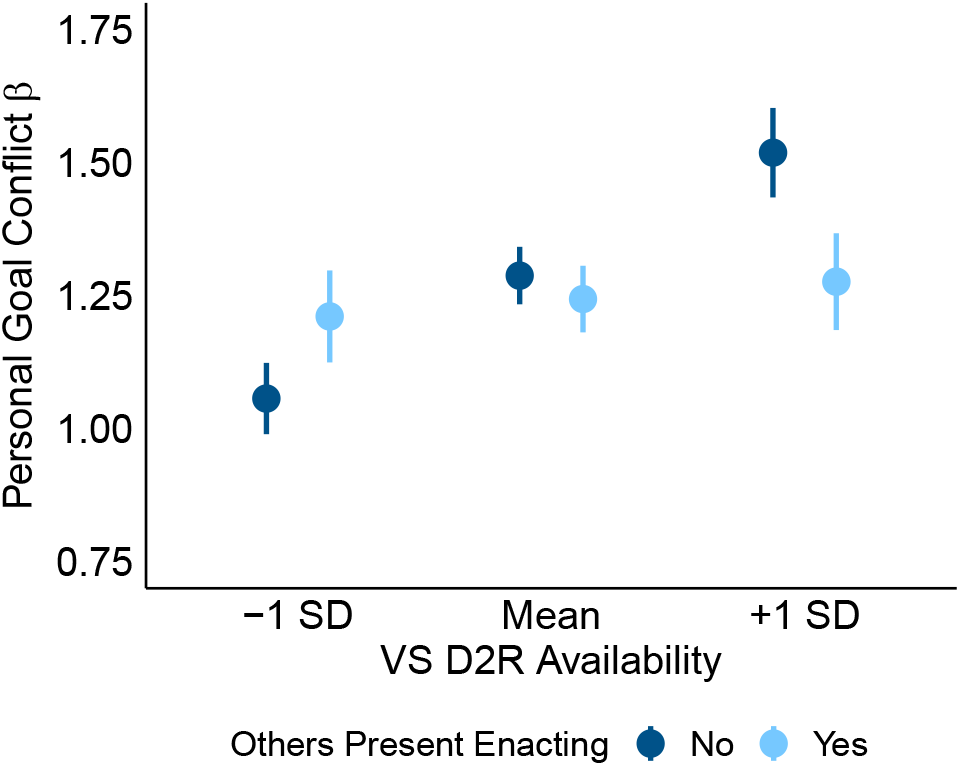
Desire resistance attempt sensitivity to personal goal conflict depends on dopamine D2 receptor availability and social context. Participants with lower dopamine D2 receptor (D2R) availability in the ventral striatum (VS) were more sensitive to their personal goal conflict when deciding whether to attempt to resist a desire while others were present engaging in that desire. Conversely, participants with higher VS D2R availability were less sensitive to their personal goal conflict in a social context than in a non-social context. Personal goal conflict sensitivities (βs) are standardized logit (log odds) regression coefficients from the simple slopes analysis. Error bars represent ± 1 standard error.

For resistance success, although there were no main effects of D2R availability in the ventral striatum (β = 0.155, CI [−0.10, 0.41], Z = 1.20, p = 0.230) or amygdala (β = 0.188, CI [−0.08, 0.46], Z = 1.38, p = 0.167), there was a main effect of midbrain D2R availability. Higher D2R availability in the midbrain was associated with better resistance success (β = 0.336, CI [0.054, 0.618], Z = 2.33, p = 0.020) (**Fig. 4a**). This effect was qualified by an interaction between midbrain D2R availability and resistance attempts (β = −0.286, CI [−0.489, −0.082], Z = - 2.76, p = 0.006) (**Fig. S1**). Simple slopes analysis indicated that higher levels of midbrain D2R availability predicted greater resistance success for unresisted desires (desires that participants did not consciously attempt to resist) (β = 0.340, CI [0.054, 0.618], Z = 2.33, p = 0.020) but not resisted desires (β = 0.050, CI [−0.226, 0.326], Z = 0.354, p = 0.724). Specifically, with each 1-SD unit increase in receptor availability, the odds of not enacting unresisted desires increased by ~0.40 (exp(0.340) = 1.41). There were no two-way interactions predicting resistance success between D2R availability in any region with either personal goal conflict or desire strength. We did not identify interactions between resistance attempt and D2R availability in the ventral striatum (β = −0.173, CI [−0.374, 0.029], Z = −1.68, p = 0.093) or amygdala (β = −0.155, CI [−0.368, 0.059], Z = −1.42, p = 0.155). Successful resistance of desires was also not moderated by the three-way interaction between degree of conflict with personal goals, resistance attempt, and D2R availability in the ventral striatum (β = 0.07, CI [−0.12, 0.26], Z = 0.743, p = 0.457), midbrain (β = −0.01, CI [−0.182, 0.158], Z = −0.138, p = 0.891), or amygdala (β = −0.19, CI [−0.383, 0.001], Z = −1.95, p = 0.051). Resistance successes were also not predicted by the threeway interaction between desire strength, resistance attempt, and D2R availability in the ventral striatum (β = −0.02, CI [−0.18, 0.14], Z = −0.269, p = 0.788), midbrain (β = −0.05, CI [−0.206, 0.107], Z = −0.619, p = 0.535), or amygdala (β = −0.06, CI [−0.236, 0.112], Z = −0.697, p = 0.486). Full model results for interactions effects of D2R availability (in each ROI) on resistance success are shown in **Table S8**.

**Fig. 4.**
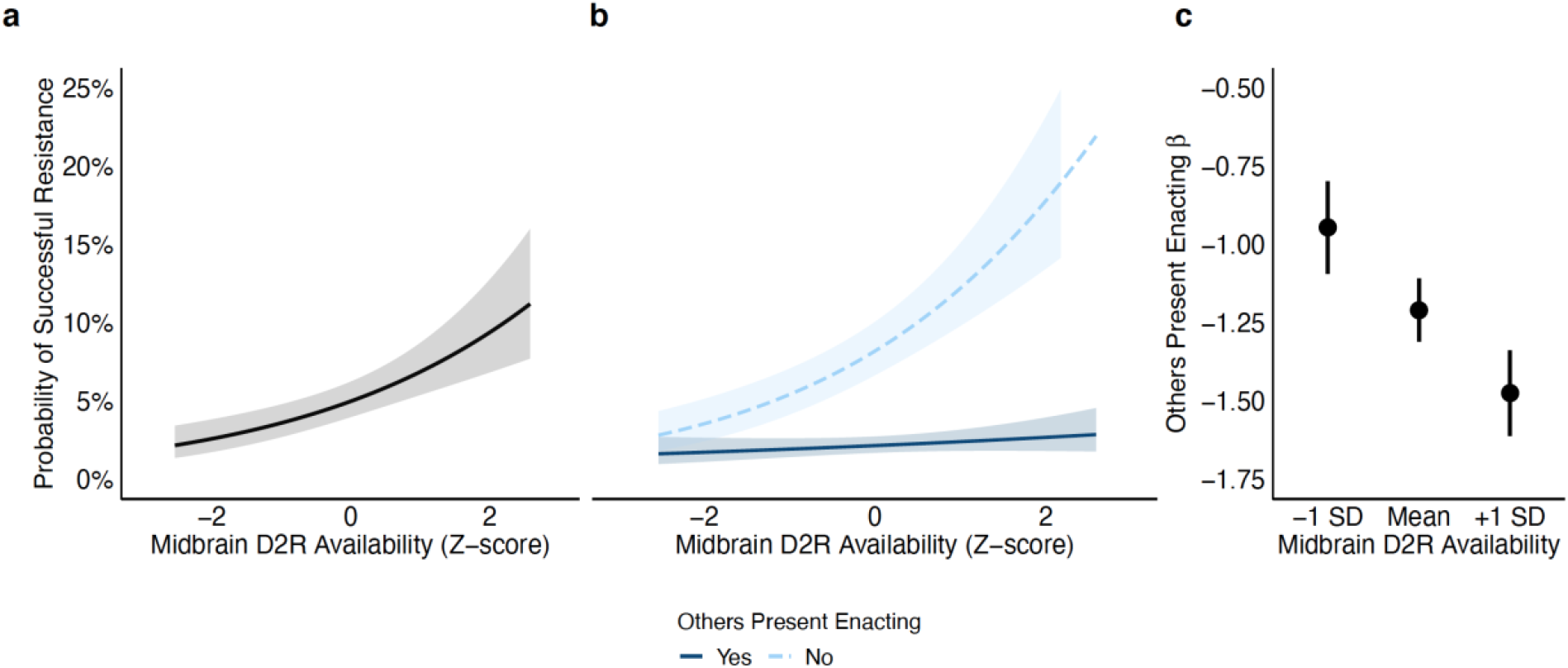
Relationship between dopamine D2 receptor availability, social context, and successful resistance of spontaneously experienced desires. Across participants, **(a)** higher midbrain D2R availability predicted a higher probability of successfully resisting a desire (attempting to resist but not enacting). Participants with higher dopamine D2 receptor (D2R) availability in the midbrain were **(b)** less successful at resist their desires in a social context (when others were present enacting the desire). Variation in the effect of social context **(c)** is further illustrated across different levels of D2R availability. Midbrain D2R availability is standardized across participants. Shaded areas and error bars represent ± 1 standard error. Social context sensitivities (βs) are standardized logit (log odds) regression coefficients from the simple slopes analysis.

To further explore these effects for successful resistance we considered whether social context influenced the effect of D2R availability. Since neither personal goal conflict nor desire strength interacted with resistance attempt to predict resistance success in the main effect models described earlier, we focused on the three-way interaction between D2R availability, resistance attempt, and social context. There was a two-way interaction between midbrain D2R availability and social context (β = −0.329, CI [−0.597, 0.536], Z = 1.06, p = 0.017) (**Fig. 4b**) but not a threeway interaction including resistance attempt (β = 0.188, CI [−0.159, −0.060], Z = −2.39, p = 0.288). Simple slopes analysis indicated differential effects of social context across levels of D2R availability. For participants with low (−1 SD) midbrain D2R availability, social context had a modest negative effect on successful resistance (β = −0.946, CI [−1.23, −0.658], Z = −6.46, p < 0.001) compared to the gradually larger negative effects for participants with average (β = −1.21, CI [−1.41, −1.01], Z = −12.0, p < 0.001) and high (+1 SD) (β = −1.47, CI [−1.74, −1.21], Z = −10.9, p < 0.001) D2R availability (**Fig. 4c**). In other words, participants with higher/+1 SD midbrain D2R availability were more sensitive to the presence of others enacting a desire (decrease in odds by ~0.77 (1-exp(−1.47) than participants with average (decrease in odds by ~0.70 (1 - exp(−1.21) or low/-1 SD (decrease in odds by ~0.61 (exp(−0.946) midbrain D2R availability. This model did not identify any significant interactions with social context and D2R availability in the ventral striatum or amygdala (**Table S10**). Full model results for interactions effects of resistance attempts, social context, and D2R availability (for each ROI) on resistance success are shown in **Table S10**.

## Discussion

In an adult life-span sample, the present study reveals that individual differences in DA function shape decisions to practice self-control over everyday desires. Individuals with higher D2R availability in the ventral striatum were more likely to consciously attempt to resist desires and were more sensitive to the level of conflict with their personal goals rather than the strength of their desires themselves in self-control decisions. Further, higher D2R availability in the midbrain, but not the ventral striatum predicted successful resistance of desires. Follow-up analyses indicated that these effects also depended on whether others were present enacting the desire. These results highlight specific mechanisms by which D2Rs bias self-control decisions.

Ventral striatum D2Rs may facilitate decisions to resist desires by amplifying the subjective benefits of conflicting personal goals over momentary temptations. In laboratory studies of humans, striatal dopamine synthesis capacity increases sensitivity to benefits versus cost of decisions involving effortful inhibition (Westbrook et al., 2020) and striatal D2Rs boosts neural representations of subjective value of delayed rewards (Castrellon et al., 2019). One interpretation is that higher ventral striatum D2R availability may provide a “boost” in control motivation that supports one’s ability to attempt to resist a desire by increasing sensitivity to the benefit of conflicting personal goals over the cost of enacting immediate desires.

We also report novel evidence that individual differences in dopamine function confers differential sensitivity to personal goal conflict on decisions to resist desires in a social context. Although on average, participants were less likely to attempt to resist a desire in the presence of others enacting that desire, those with high ventral striatal D2R availability showed a greater sensitivity to goal conflict when alone but this enhanced sensitivity was largely eliminated in the presence of others enacting the desire. By contrast, those with low ventral striatal D2R not only showed no detriment in their sensitivity to goal conflict but appeared to show enhanced sensitivity when others were enacting the desire. This finding is intriguing in that it suggests a neuropharmacological feature may influence the extent to which social influences, such as conformity pressures, influence one’s balancing of conflicting personal goals with immediate desires. While having higher D2R availability may facilitate decisions to resist temptations particularly when those desires strongly conflict with goals, that sensitivity to goal conflict appears to be eliminated by social pressures. While this specific pattern was not predicted, it is consistent with the sensitivity of the ventral striatum to social variables. fMRI studies show that activation in the ventral striatum is heightened when people receive rewards that were known to be valued by others (Campbell-Meiklejohn et al., 2010) and activity in this region is sensitive to conflict or mismatch with value ratings from other people (Klucharev et al., 2009). The ventral striatum also exhibits heightened activation during the anticipation of affiliative social rewards (Bortolini et al., 2021). Ventral striatal dopamine function could therefore support preferences for conformity by downweighing the value of conflicting long-term personal goals to better align with behaviors of others present. Conversely, this could also suggest that low D2R participants do not find social conformity or matching behavior to be rewarding. An alternative explanation is that individuals with lower D2R availability value conflicting personal goals more because observing others enact the desire increases their own perceived self-efficacy via aspiration to perform well. This explanation is in line with results from a candidate gene study showing that a single nucleotide polymorphism associated with lower D2R affinity (C957T) is related to higher levels of social facilitation (Walter et al., 2011). Social facilitation has been linked to better selfcontrol among individuals high in impression management (a personality profile linked to low dominance and high affiliation) (Uziel & Baumeister, 2012). This explanation is less straightforward, but given that low D2R availability has been correlated with higher submissiveness (Cervenka et al., 2010) and attachment (Caravaggio et al., 2017), future work should seek to evaluate how dopamine interacts with traits and social motivations to conform.

Unlike DA receptors in the striatum, midbrain D2Rs largely function as somatodendritic autoreceptors that downregulate dopamine cell firing and release (Ford, 2014). Consistent in direction with prior associations with impulsive traits (Buckholtz et al., 2010; Zald et al., 2008), here individuals with higher autoreceptor availability were more likely to be successful in preventing desired acts. This effect was modest, and interestingly appeared to primarily reflect a lower enactment of desires that were not consciously resisted (i.e., in the absence of a resistance attempt). Given that autoreceptor levels did not significantly impact desire enactment when there was a conscious attempt to resist the desires, the autoreceptor impact on behavior may be speculated to reflect a relatively subtle automatic effect that is less important in the context of more conscious efforts to resist desires. The influence of midbrain autoreceptors did not interact with underlying motivations like personal goal conflict or desire strength. However, its influence interacted with whether the desire was experienced in the presence of others enacting the desire. While on average, participants were more likely to enact a desire when it was experienced in the presence of others enacting that desire, participants with higher midbrain D2R availability were more sensitive to this social context than participants with low autoreceptor availability. Thus, whatever automatic benefit having higher midbrain autoreceptors has in successful desire resistance, it is largely eliminated in the face of social influence and conformity pressures. These interactions point to a role for affiliation goals and social cognition in determining context sensitivity of the dopamine system. For example, evidence in humans shows that fMRI activation in the midbrain related to social reward cues is heightened in people who crave social interaction following social isolation (Tomova et al., 2020). It is worth noting that although we asked participants to indicate whether others are present enacting the desire they are experiencing, social influences may be quite different in the presence of others who are not enacting a desire. Future work should explore how dopaminergic variables influence responses to different types of social context. Although the present study was restricted to adults it is interesting to speculate how these interactions may play out in adolescence given current thinking about dopamine’s specific role in adolescent behavior and the importance of peer pressure and conformity pressures in this developmental period (Telzer, 2016).

In addition to the identification of specific neurobiological mechanisms of self-control, the findings also make more general theoretical contributions to our understanding of self-control. The observation that VS DA biases attempts to resist desires and that midbrain DA biases enactment of desires could suggest that these regions have distinct influence over different aspects of self-control. When participants initially experience a desire, they engage meta-cognitive processes that help evaluate whether self-control is possible or even necessary (e.g. when there is only weak conflict between desire strength and personal goals) (Boureau et al., 2015). This kind of decision has been previously characterized as a judgment about control motivation and influences the amount of potential control effort to expend by attempting to resisting a desire (Kotabe & Hofmann, 2015). With sufficient control motivation and potential control effort, a person can generate enough actual control effort in their attempt to resist a desire. Then, depending on the amount of actual control expended, individuals either enact or resist their desire. Our observation that social context is a critical determinant of desire resistance is in line with evidence that social influence affects effort expenditure for rewards (Gilman et al., 2015).

Given the importance of ventral striatal dopamine in wanting processes, and their relevance for addiction (Berridge & Robinson, 2016), it is notable that D2R availability did not alter the extent to which desire strength determines resistance attempts. Adding to this, exploratory correlation analyses indicated that having more or fewer D2Rs did not relate to the frequency or strength of spontaneous desires (**Table S2**). This aligns with evidence that dopamine does not globally alter sensitivity to rewards per se but is more specific to rewards with an acquisition cost (Hamid et al., 2016; Phillips et al., 2007; Salamone & Correa, 2012). For instance, when rats are depleted of dopamine, their overall consumption of food remains unaltered; instead, dopamine affects the weighting of responses costs associated with lever pressing to obtain food (Cousins & Salamone, 1994; Salamone et al., 1991). Similarly, pharmacological studies in rats and humans indicate that dopamine effects on reward processes emerge when effort costs increase (Floresco et al., 2008; Wardle et al., 2011). The present results may thus be interpreted in a similar light, but with the cognitive effort to resist the urge, for the sake of achieving a long-term goal providing the key point of ventral striatal D2R influence. Future studies in which participants also rate the subjective effort involved in resisting urges will be needed to test this speculation.

Here we asked participants about the strength of their desires and the degree to which those desires conflict with personal goals. Future studies can test competing models of self-control (Berkman et al., 2017; Shenhav, 2017) with additional experience sampling questions that isolate features like desire attributes, choice difficulty, perceived level of effort needed to resist an urge, cognitive resource allocation, desire proximity or accessibility, goal congruency, and personal identity since these factors can influence self-control (Berkman et al., 2017; Frömer et al., 2019) and dopamine-mediated differences in subjective value (Westbrook et al., 2020). Further, while we asked participants to identify whether others were around enacting the desire, experimental designs could probe whether social conformity effects operate in the opposite direction by asking whether others are present resisting a desire since affiliation could support enhanced self-control (Cullum et al., 2011). The observed context-dependent effects of dopamine also offer insight into potential individualized self-control clinical interventions. For example, making efforts to change ones social situation can help reduce the potential for harmful social cues to affect drug use, eating, or exercise (Duckworth et al., 2016, 2018). The efficacy of these social situation change strategies may be more effective for participants depending on their reward sensitivity to long-term goals and dopamine function. Given that substance use disorders have been linked to lower dopamine D2R availability (Ashok et al., 2017, 2019), interventions could explore the impact of social influence on how these populations weigh long-term benefits of drug resistance.

An unavoidable drawback of neuroscience research is that observed behavior in controlled lab tasks may not always reflect the natural and highly idiosyncratic behaviors that individuals express in their day-to-day lives. Using experience sampling, our findings overcome this challenge in the context of neurotransmitter function and desire resistance and contribute novel insight to our understanding of generalizable neural markers of self-control in everyday life. Controlled laboratory experiments remain invaluable and complementary to experience sampling methods because they can explain behavior under common contexts. Future investigations can benefit from integrating approaches that approximate behavior in both stable and variable environments. While the lack of a common context for observing behavior has its own limitations, the extension to self-control behavior in everyday life also emphasizes the generality with which dopaminergic variables help support self-control across a wide range of common desire types and personal goals. Such associations demonstrate the utility of this multi-method approach to further understand neural mechanisms of other spontaneous self-control decisions such as those involving consumer behaviors, compulsive drug use, and social norm-conformity violations.

## Acknowledgements

This research was supported by National Institute on Aging grants R01-AG044838 and R01-AG043458, and National Institute on Drug Abuse grant R21-DA033611. J.J.C. was supported by an NSF Graduate Research Fellowship (grant no. NSF DGE-1644868).

## Data

Data and code used in the manuscript can be viewed and downloaded from OSF: https://osf.io/wa36m/

---

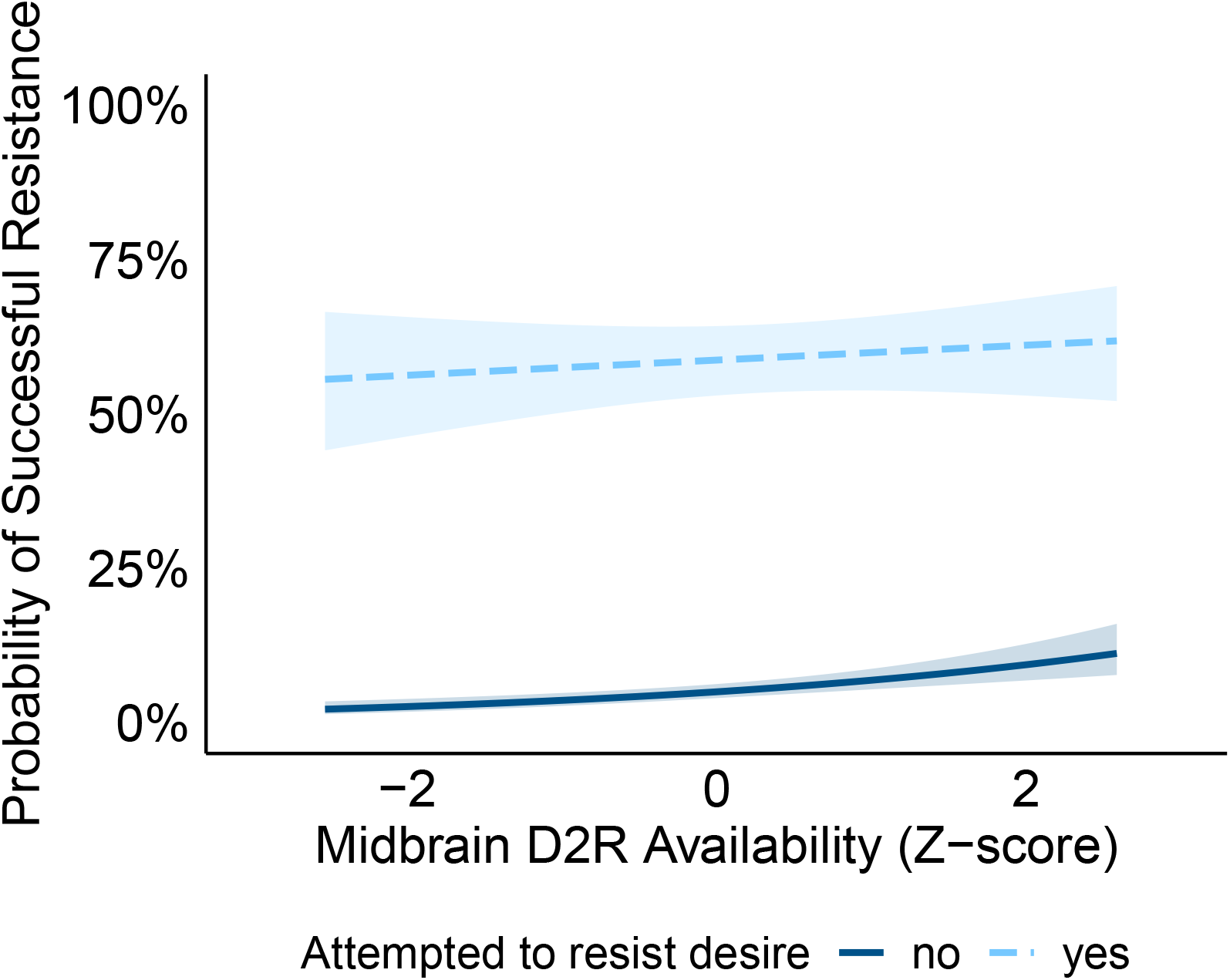

## Notes

https://osf.io/wa36m/

### Competing Interest Statement

The authors have declared no competing interest.

