## Supplemental materials for "Dopamine biases sensitivity to personal goals and social influence in self-control over everyday desires"

### Supplementary Materials

*Participants and Procedures*

The data analyzed here were collected from three study samples. They are described as samples 1, 2, and 3. Although the overall focus of each study was different, participants in each sample completed the same PET imaging and experience sampling procedures – which produced the data analyzed in the present manuscript. Sample 1 included 20 young adults (ages 18–24, M = 21.1, SD = 1.92, 10 females) recruited from the Vanderbilt University community in Nashville, TN between 2012 and 2013. Sample 2 included 40 healthy adults (ages 22–80, M = 44.7, SD = 17.6, 21 females) recruited from the Greater Nashville, TN metropolitan area between 2013 and 2016. Sample 3 included 43 healthy adults (ages 20–65, M = 42.0, SD = 15.4, 22 females) recruited from the Greater Nashville, TN metropolitan area between 2016 and 2018. Data from all samples were collected at Vanderbilt University **(See Table S1 for descriptive statistics for each sample)**. In each sample, participants were subject to the following exclusion criteria: any history of psychiatric illness on a screening interview (a Structural Interview for Clinical DSM-IV Diagnosis was also available for all subjects and confirmed no history of major Axis I disorders) (First et al., 1997), any history of head trauma, any significant medical condition, or any condition that would interfere with MRI (e.g. inability to fit in the scanner, claustrophobia, cochlear implant, metal fragments in eyes, cardiac pacemaker, neural stimulator, pregnancy, and metallic body inclusions or other contraindicated metal implanted in the body). Participants with major medical disorders including diabetes and/or abnormalities on a comprehensive metabolic panel, complete blood count, or EKG were excluded. Participants were also excluded if they reported a history of substance abuse, current tobacco use, alcohol consumption greater than 8 ounces of whiskey or equivalent per week, use of psychostimulants (excluding caffeine) more than twice at any time in their life or at all in the past 6 months, or any psychotropic medication in the last 6 months other than occasional use of benzodiazepines for sleep. Any illicit drug use in the last 2 months was grounds for exclusion, even in participants who did not otherwise meet criteria for substance abuse. Urine drug tests were administered, and subjects testing positive for the presence of amphetamines, cocaine, marijuana, PCP, opiates, benzodiazepines, or barbiturates were excluded. Pre-menopausal females had negative pregnancy tests at intake and on the day of the scan.

*Positron Emission Tomography Imaging*

[18F]fallypride, (S)-N-[(1-allyl-2-pyrrolidinyl)methyl]-5-(3[18F]fluoropropyl)-2,3-dimethoxybenzamide was produced in the radiochemistry laboratory attached to the PET unit at Vanderbilt University Medical Center, following synthesis and quality control procedures described in US Food and Drug Administration IND 47,245. PET data were collected on a GE Discovery STE (DSTE) PET scanner (General Electric Healthcare, Chicago, IL, USA). Serial scan acquisition was started simultaneously with a 5.0 mCi slow bolus injection of DA D2/3 tracer [18F]fallypride (specific activity greater than 3000 Ci/mmol). CT scans were collected for attenuation correction prior to each of the three emission scans, which together lasted approximately 3.5 h with two breaks for subject comfort. Prior to the PET scan, T1-weighted magnetic resonance (MR) images (TFE SENSE protocol; Act. TR = 8.9 ms, TE = 4.6 ms, 192 TFE shots, TFE duration = 1201.9 s, FOV = 256 × 256 mm, voxel size =1 × 1 × 1 mm) were acquired on a 3T Philips Intera Achieva whole-body scanner (Philips Healthcare, Best, The Netherlands). In Study 1 and 2, subjects were scanned only once. In study 3, subjects were scanned as part of a blinded two scan protocol, in which the one scan analyzed here was conducted 3-hours after placebo administration, while an additional scan (not analyzed here) was conducted following administration of oral amphetamine. Approval for the study protocol was obtained from the Vanderbilt University Human Research Protection Program and the Radioactive Drug Research Committee. All participants in each sample completed written informed consent. Each samples' study procedures were approved in accordance with the Declaration of Helsinki's guidelines for the ethical treatment of human participants.

*Binding Potential Calculation*

We estimated D2 receptor availability as binding potential ($BP_{ND}$) using the simplified reference tissue model (SRTM) with the cerebellum as the reference region(Lammertsma & Hume, 1996). The midbrain was drawn in MNI standard space using previously described guidelines (Dang, Donde, et al., 2012; Dang, O'Neil, et al., 2012; Mawlawi et al., 2001) and registered to PET images using the same transformations used in $BP_{ND}$ calculation. All other ROIs were derived from the Hammers Atlas plus deep nuclei parcellation as produced from the parcellation of the T1 structural image of each subject in the PNEURO module of PMOD software. The PET data were registered to the T1 image for each subject and, thus, to the ROIs (all steps implemented in PNEURO module of PMOD Software). $BP_{ND}$ values from ROIs were obtained by fitting the SRTM to the PET time activity curve data from each ROI in the PKIN (kinetic modeling) module of PMOD using the cerebellum as the reference region. These ROI-based $BP_{ND}$ values were then averaged across hemispheres. Our lab and others have shown that many brain regions may be susceptible to partial volume effects in estimating $BP_{ND}$ especially in older adults as a result of age differences in gray matter volume (Smith et al., 2019). Therefore, we used PVC values in all analyses presented here with the exception of the midbrain for which we used uncorrected $BP_{ND}$ for analysis, because it was not available in the Hammers Atlas in PNEURO.

*Experience Sampling*

During a visit to the lab as part of larger multi-day studies, participants were first asked to provide their typical times for: waking, sleeping, eating lunch, and eating dinner. These time-points served as individually-tailored boundaries for text message deliveries to ensure that messages were not disruptive of participants' sleep. Text messages with links to the web-based survey were programmed for delivery using OhDontForget.com. Each participant received three text messages per day for ten days (90 messages total). The exact time of delivery per day was pseudorandomized based on the time points of waking, sleeping, and eating provided by the participant. The experience sampling protocol was adapted from (Hofmann, Baumeister, et al., 2012; Hofmann, Vohs, et al., 2012).

Participants were instructed to respond to the messages (by clicking on the link and completing the survey in the web browser on their phone) as soon as they received them as long as they could safely do so (e.g. if participants received a text message while driving, they were instructed to wait until their drive was complete/when it was safe to respond). If participants did not have a smartphone that could render the survey webpage, a smartphone was provided for them to use during the course of ESM data collection. Participants were instructed to report their strongest desires experienced in the last three hours. Each survey permitted entry of up to three desires.

Participants were asked to indicate where they were and if they experienced a desire (craving, urge, longing) in the past three hours. They were provided with 13 desire options:

1. Eating, snacking, nonalcoholic drinks
2. Alcohol, cigarettes, tobacco, other drugs
3. Entertainment media (TV, movies, web browsing, video games)
4. Social Media (Facebook, Twitter, Instagram, etc.)
5. Spending
6. Sex
7. Sleep
8. Social Contact (in person or phone conversation, texting, FaceTime, etc.)
9. Leisure and relaxation
10. Exercise
11. Work
12. Other
13. None

If participants selected "Other," they were prompted to provide a description of their desire. After selecting a desire, participants indicated the strength of the desire on a scale from 0 (no desire at all) to 7 (irresistible). Next, participants were asked how much the desire conflicted with other personal goals on a scale from 0 (no conflict at all) to 4 (very high conflict). If they identified that the desire conflicted with another goal, the participants were asked to categorize the conflicting goal. They were provided with 12 goal options:

1. Healthy living (healthy eating, healthy drinking, reducing health damage, bodily fitness, good appearance, ending dependency)
2. Saving money
3. Abstinence
4. Educational achievement
5. Professional achievement
6. Sport achievement
7. Social goals (social appearance, social recognition, socializing)
8. Moral Integrity (fidelity, religious/cultural beliefs)
9. Using time efficiently
10. Not delaying things and getting things done
11. Leisure and Relaxation
12. Other

If participants selected "Other" they were once again prompted to provide a description of their desire. Finally, the participants were asked to indicate if they attempted to resist the desire (Yes or No), if they enacted the desire (Yes or No), and if other people present (either physically or via media) were enacting the desire (Yes or No).

The most frequently reported desires were "Eating, snacking, nonalcoholic drinks," "Sleep," and "Leisure and Travel." Desires reported as "other" included activities that ranged from playing music and writing to chores like cleaning or showering.

*Statistical Analysis*

Prior to advanced modelling, we conducted simple correlation analyses between D2R availability and participants' summarized untransformed ratings and decisions. Specifically, we examined correlations between D2R availability (in each of 3 ROIs: ventral striatum, midbrain, and amygdala) and: (1) total number of desires experienced, (2) total number of attempts to resist, (3) total number of desires enacted, (4) proportion of attempts made (out of experienced), (5) proportion of desires enacted (out of experienced), (6) average desire strength rating, and (7) average personal goal conflict rating. The results of these correlations are reported in **Table S2**. D2R availability was not correlated for any ROI with the total number of desires reported, providing some assurance that DA did not relate to survey compliance. From all these analyses across all ROIs, only 3 statistically significant effects were identified with midbrain D2R availability: (1) total number of desires enacted ($r = -0.264$, $p = 0.007$), (2) proportion of desires resisted ($r = 0.273$, $p = 0.005$), and (3) average personal goal conflict rating ($r = 0.203$, $p = 0.039$). Although these effects are consistent in direction with expected role of D2R availability in downregulation of dopamine release and higher self-control, these effects are fragile and did not remain statistically significant after controlling for age, sex, and study sample, and number of days between the PET scan day and the first day of experience sampling as covariates in linear regression tests: (1) total number of desires enacted ($t(97) = -1.70$, $\beta = -3.67$, $p = 0.091$), (2) proportion of attempts to resist ($t(97) = 0.414$, $\beta = 0.010$, $p = 0.680$), and (3) average personal goal conflict rating ($t(97) = -0.230$, $\beta = -0.065$, $p = 0.819$).

We fit each participant's ratings to several different models that differ in how personal goals and desires are integrated to support decisions to attempt to resist desires. We considered the possibility that (1) desires are linearly discounted from personal goals, (2) personal goals are normalized by desire strength, or that (3) personal goals and desire strength independently contribute to resistance attempts. Since participants varied in the number of desires reported and in the range of ratings reported (which included zero), we added 1 to all ratings and standardized ratings within each participant's set of responses.

The data analyzed only includes participants who responded to at least half of the minimum possible number of surveys (15/30 minimum surveys). Data also excludes participants who did not vary at all in either their goal conflict or desire strength ratings. Prior to analysis, we excluded extreme desire and goal conflict ratings with a within-participant standardized Z-score +/- 3. This resulted in exclusion of 69 individual desires. These exclusions did not amount to exclusion of any participants. For transparency, we also present the results with these extreme values, here, to demonstrate that our effects are extremely robust to outlier exclusion. This provided a final sample size of $N = 103$. All descriptive statistics (including those indicated in the Participants and Procedures section above) reflect data from these final 103 participants.

To account for within- and between-participant differences, we carried out multilevel models using mixed-effects analysis in R using the lme4 package and fit using the glmer function with the BOBYQA (Bound Optimization BY Quadratic Approximation) (Powell, 2009) optimizer and 200,000 functional evaluations. All models included random intercepts for participants. To ensure primary results could not be accounted for by confounds of age, sex, or study sample, these three variables were included in primary analyses as fixed effects. This was particularly important for age given its known impact on both D2R as well as evidence that age impacts self-control over everyday desires (Burr et al., 2020; Karrer et al., 2017). In addition to ratings, we standardized ages and D2R availability across across participants in the entire dataset. There were differences across the samples in average $BP_{ND}$ in all regions of interest (Table S1). However, additional analyses suggested that adding random intercepts for study sample was not necessary (intraclass coefficients between <.0001 and .013 across all models) and inclusion had a negligible impact on model coefficients.

Models included logistic regressions for binary outcomes of attempts to resist desires or failed attempts to resist desires. We initially fit the following models in equations 1-5 below to identify main effects of personal goals and/or desires. Results from these models are in **Table S3**. We evaluated model fit quality using Bayesian Information Criterion (BIC) scores. The model in equation 5 had the lowest BIC score, indicating that it provided the overall best fit. This model implies that personal goals and desires independently contribute to self-control decisions.

$$\Pr(y_{ij}) = logit^{-1}\big(\beta_0 + \beta_1(Goal\,)_{ij} + \beta_2(Age)_j + \beta_3(Sex)_j + \beta_4(Sample)_j\big) + u_{0j} + e_{0ij} \qquad \textit{Eq. 1}$$

$$\Pr(y_{ij}) = logit^{-1}\big(\beta_0 + \beta_1(Desire\,)_{ij} + \beta_2(Age)_j + \beta_3(Sex)_j + \beta_4(Sample)_j\big) + u_{0j} + e_{0ij} \qquad \textit{Eq. 2}$$

$$\Pr(y_{ij}) = logit^{-1}\big(\beta_0 + \beta_1(Goal\, - \,Desire)_{ij} + \beta_2(Age)_j + \beta_3(Sex)_j + \beta_4(Sample)_j\big) + u_{0j} + e_{0ij} \qquad \textit{Eq. 3}$$

$$\Pr(y_{ij}) = logit^{-1}\big(\beta_0 + \beta_1(Goal\,/Desire)_{ij} + \beta_2(Age)_j + \beta_3(Sex)_j + \beta_4(Sample)_j\big) + u_{0j} + e_{0ij} \qquad \textit{Eq. 4}$$

$$\Pr(y_{ij}) = logit^{-1}\big(\beta_0 + \beta_1(Goal\,)_{ij} + \beta_2(Desire\,)_{ij} + \beta_3(Age)_j + \beta_4(Sex)_j + \beta_5(Sample)_j\big) + u_{0j} + e_{0ij} \qquad \textit{Eq. 5}$$

Where $i =$ survey-level effects and $j =$ participant-level effects with variances defined as:
$e_{0ij} \sim N(0, \sigma_e^2)$ for fixed effects and
$\mu_{0j} \sim N(0, \sigma_\mu^2)$ for random effects

and resistance attempts are coded as:

$$y_{ij}\begin{cases} 1\; if\; y_{ij} = Attempt \\ 0\; if\; y_{ij} = No\ Attempt \end{cases}$$

Desire enactment was modeled in equation 6 as the interaction between situational variables and attempts to resist (dummy-coded as 1 for attempt and 0 for no attempt). We note that BIC values for the model defined in equation 6 and one using a single difference term (personal goal conflict minus desire strength) were relatively similar (compared to an intercept model): two-parameter goal, desire model BIC = 4311.26 versus single parameter goal minus desire model BIC = 4296.08. Although the single-parameter model was a slightly better fit to the data, both models explained the same amount of variance (fixed-effects $R^2 = 0.44$, total $R^2 = 0.57$). Therefore, to maintain consistency with the resistance attempt model in equation 5, we use the two-parameter model defined below in equation 6. Full model results from this model (and comparison to other models as above for resistance attempts) are in **Table S4**.

$$\Pr(y_{ij}) = logit^{-1}\big(\beta_0 + \beta_1(Goal\,)_{ij} + \beta_2(Desire\,)_{ij} + \beta_3(Attempt\,)_{ij} + \beta_4(Goal)_{ij} * (Attempt)_{ij} + \beta_5(Desire)_{ij} * (Attempt)_{ij} + \beta_6(Age)_j + \beta_7(Sex)_j + \beta_8(Sample)_j\big) + u_{0j} + e_{0ij} \qquad \textit{Eq. 6}$$

where resistance success is coded as:

$$y_{ij}\begin{cases}1\ if\ y_{ij} =\ No\ Enact\\ 0\ if\ y_{ij} = Enact\end{cases}$$

Interactions with D2R availability were fit according to the model in equations 5 and 6 to evaluate independent effects of D2R availability on goals and desires in equations 7 and 8, below, to identify relationships with predicted resistance attempts and successful resistance. Results from these models are in **Tables S5** (resistance attempt) and **Table S6** (resistance success).

Resistance attempt:

$$\begin{aligned}\Pr(y_{ij}) = logit^{-1}\big(&\beta_0 + \beta_1(Goal\ )_{ij} + \beta_2(Desire\ )_{ij} + \beta_3(Age)_j\\ &+\ \beta_4(Sex)_j + \beta_5(Sample)_j + \beta_6(D2R_{avail})_j\\ &+ \beta_7(PET - EMA_{date\ \Delta})_j + \beta_8(Goal)_{ij} * (D2R_{avail})_j\\ &+ \beta_9(Desire)_{ij} * (D2R_{avail})_j\big) + u_{0j} + e_{0ij}\end{aligned} \qquad Eq.\ 7$$

Successful resistance:

$$\begin{aligned}\Pr(y_{ij}) = logit^{-1}\big(&\beta_0 + \beta_1(Goal\ )_{ij} + \beta_2(Desire\ )_{ij} + \beta_3(Attempt\ )_{ij}\\ &+ \beta_4(Age)_j +\ \beta_5(Sex)_j + \beta_6(Sample)_j + \beta_7(D2R_{avail})_j\\ &+ \beta_8(PET - EMA_{date\ \Delta})_j + \beta_9(Goal)_{ij} * (Attempt)_{ij}\\ &+ \beta_{10}(Attempt)_{ij} * (D2R_{avail})_j + \beta_{11}(Goal)_{ij}(D2R_{avail})_j\\ &+ \beta_{12}(Goal)_{ij} * (Attempt)_{ij} * (D2R_{avail})_j + \beta_{13}(Desire)_{ij}\\ &* (Attempt)_{ij} + \beta_{14}(Desire)_{ij}(D2R_{avail})_j + \beta_{15}(Desire)_{ij}\\ &* (Attempt)_{ij} * (D2R_{avail})_j\big) + u_{0j} + e_{0ij}\end{aligned} \qquad Eq.\ 8$$

To facilitate interpretation of cross-level interactions, we conducted simple slopes analysis to identify ORs for participants with low (-1 SD), average, and high (+1 SD) D2R availability. Simple slopes analysis was carried out using the “jtools” and “interactions” R packages.

For models evaluating social context effects, a linear mixed-effects models in equation 9 and equation 10 tested the interaction between the presence of others enacting the desire and desire ratings (for resistance attempt) and the interaction with resistance attempt (for resistance success). Results from these models are in **Table S9 and Table S10**. The interaction between midbrain D2R availability and resistance attempt predicting resistance success is plotted in **Figure S1**. Resistance attempt was dummy-coded as 1 for attempt and 0 for no attempt and presence of others enacting was dummy-coded as 1 for yes and 0 for no.

Resistance attempt:

$$\begin{aligned}\Pr(y_{ij}) = logit^{-1}\big(&\beta_0 + \beta_1(Goal\ )_{ij} + \beta_2(Desire\ )_{ij}\\ &+ \beta_3(Others\ Present\ \ Enacting)_{ij} + \beta_4(Age)_j +\ \beta_5(Sex)_j\\ &+ \beta_6(Sample)_j\big) + \beta_7(Goal\ )_{ij}\\ &* (Others\ Present\ Enacting\ )_{ij} + \beta_8(Desire\ )_{ij}\\ &* (Others\ Present\ Enacting)_{ij} + u_{0j} + e_{0ij}\end{aligned} \qquad Eq.\ 9$$

where resistance attempt is coded as:

$$y_{ij}\begin{cases} 1\ if\ y_{ij} =\ Attempt \\ 0\ if\ y_{ij} = No\ Attempt \end{cases}$$

Resistance success:

$$\begin{aligned}\Pr(y_{ij}) = logit^{-1}\big(&\beta_0 + \beta_1(Goal\ )_{ij} + \beta_2(Desire\ )_{ij} +\ \beta_3(Attempt\ )_{ij} \\ &+ \beta_4(Others\ Present\ \ Enacting)_{ij} + \beta_5(Age)_j +\ \beta_6(Sex)_j \\ &+ \beta_7(Sample)_j\big) + \beta_8(Goal\ )_{ij} \\ &* (Others\ Present\ Enacting\ )_{ij} + \beta_9(Desire\ )_{ij} \\ &* (Others\ Present\ Enacting)_{ij} + \beta_{10}(Attempt\ )_{ij} \\ &* (Others\ Present\ Enacting)_{ij} + \beta_{11}(Goal\ )_{ij} \\ &* (Others\ Present\ Enacting)_{ij} * (Attempt)_{ij} \\ &+ \beta_{11}(Desire\ )_{ij} * (Others\ Present\ Enacting)_{ij} \\ &* (Attempt)_{ij} + u_{0j} + e_{0ij}\end{aligned} \qquad \textit{Eq. 10}$$

where resistance success is coded as:

$$y_{ij}\begin{cases} 1\ if\ y_{ij} =\ No\ Enact \\ 0\ if\ y_{ij} = Enact \end{cases}$$

Moderating effects of social context on D2R availability in resistance attempt decisions were assessed using three-way interactions in equation 11 below.

$$\begin{aligned}\Pr(y_{ij}) = logit^{-1}\big(&\beta_0 + \beta_1(Goal\ )_{ij} + \beta_2(Desire\ )_{ij} \\ &+ \beta_3(Others\ Present\ \ Enacting)_{ij} + \beta_4(Age)_j +\ \beta_5(Sex)_j \\ &+ \beta_6(Sample)_j\big) + \beta_7(D2R_{avail})_j + \beta_8(PET - EMA_{date\ \Delta})_j \\ &+ \beta_9(Goal\ )_{ij} * (Others\ Present\ Enacting\ )_{ij} \\ &+ \beta_{10}(Desire\ )_{ij} * (Others\ Present\ Enacting)_{ij} \\ &+ \beta_{11}(Others\ Present\ Enacting\ )_{ij} * (D2R_{avail})_j \\ &+ \beta_{12}(Goal\ )_{ij} * (Others\ Present\ Enacting\ )_{ij} * (D2R_{avail})_j \\ &+ \beta_{13}(Desire\ )_{ij} * (Others\ Present\ Enacting)_{ij} \\ &* (D2R_{avail})_j + u_{0j} + e_{0ij}\end{aligned} \qquad \textit{Eq. 11}$$

where resistance attempt is coded as:

$$y_{ij}\begin{cases} 1\ if\ y_{ij} =\ Attempt \\ 0\ if\ y_{ij} = No\ Attempt \end{cases}$$

Since neither personal goal conflict nor desire strength interacted with resistance attempt to predict resistance success in the main effect models described earlier, we focused on the three-way interaction between D2R availability, resistance attempt, and social context in equation 12 below.

$$\begin{aligned}\Pr(y_{ij}) = logit^{-1}\big(\beta_0 + \ &\beta_1(Attempt\ )_{ij} + \beta_2(Others\ Present\ \ Enacting)_{ij} \\ &+ \beta_3(Age)_j + \ \beta_4(Sex)_j + \beta_5(Sample)_j\big) + \beta_6(D2R_{avail})_j \\ &+ \beta_7(PET - EMA_{date\ \Delta})_j + \beta_8(Attempt\ )_{ij} \\ &* (Others\ Present\ Enacting\ )_{ij} + \beta_9(Attempt\ )_{ij} * \\ &* (\ D2R_{avail})_j + \beta_{10}(Others\ Present\ Enacting\ )_{ij} \\ &* (\ D2R_{avail})_j + \beta_{11}(Attempt\ )_{ij} \\ &* (Others\ Present\ Enacting\ )_{ij} * (\ D2R_{avail})_j + u_{0j} + e_{0ij}\end{aligned} \qquad \textit{Eq. 12}$$

where resistance success is coded as:

$$y_{ij}\begin{cases}1\ if\ y_{ij} = \ No\ Enact \\ \ 0\ if\ y_{ij} = Enact\end{cases}$$

*Follow-up Models: Inclusion of extreme outlier observations*

As described above, we limited analysis to EMA variables that fell within a standardized Z-score +/- 3. We provide the coefficients tables which demonstrate that are results are largely unchanged, and, in some cases, strengthened. We report resistance attempt and resistance success in **Table 11** and **Table 12**, respectively. We report interactions with D2R availability for each ROI for resistance attempt and resistance success in **Table 13** and **Table 14**, respectively. For the social context model, we report the interaction between desire ratings and presence of others enacting for the resistance attempt model and the interaction between the presence of others enacting and resistance attempt for the resistance success model in **Tables 15** and **Table 16**. Finally, we report the three-way interaction between D2R availability, social context, and desire ratings for the resistance attempt model and the three-way interaction between D2R availability, social context, and resistance attempt for the resistance success model in **Tables 17** and **Table 18**. Notably, the interaction between amygdala D2R availability and personal goal conflict predicting resistance attempt was statistically significant after including outliers (β = 0.098, CI [0.016, 0.180], Z = 2.34, p = 0.020). The interaction between midbrain D2R availability and personal goal conflict predicting resistance attempt was also statistically significant after including outliers (β = 0.085, CI [0.005, 0.166], Z = 2.07, p = 0.038). For social context models, the three-way interaction between presence of others enacting, personal goal conflict, and D2R availability in the amygdala was statistically significant (β = -0.21, CI [-0.37, -0.04], Z = -2.47, p = 0.014), exhibiting the same pattern reported for the ventral striatum. For successful resistance, the interaction between presence of others enacting and D2R availability in the ventral striatum was also statistically significant (β = 0.28, CI [0.00, 0.57], Z = 1.97, p = 0.049), exhibiting the opposite pattern reported for the midbrain.

### Supplementary Tables and Figures

### Table S1. Participant demographics across samples.

| | Sample 1 | Sample 2 | Sample 3 | |
|---|---|---|---|---|
| N | 20 | 40 | 43 | - |
| Age | 21.1 ± 1.92 | 44.7 ± 17.6 | 42.0 ± 15.4 | $F(2,100) = 18.2$, $p < .001$ |
| Sex | 10 F, 10 M | 21 F, 19 M | 22 F, 21 M | $X^2$ (2, N=103) = .036, $p = .982$ |
| Years Education | 14.8 ± 1.43 | 16.2 ± 1.99 | 16.0 ± 2.45 | $F(2,98) = 3.13$, $p = .048$ |
| Mean Household Income | - | $60K – 70K | $60K – 70K | $F(1,81) = .511$, $p = .477$ |
| Midbrain $BP_{ND}$ | 1.52 ± .260 | 1.22 ± .241 | 1.44 ± .304 | $F(2,100) = 10.3$, $p < .001$ |
| Ventral Striatum $BP_{ND}$ | 32.3 ± 9.65 | 39.4 ± 8.41 | 33.5 ± 8.89 | $F(2,100) = 6.20$, $p = .002$ |
| Amygdala $BP_{ND}$ | 2.97 ± .590 | 3.25 ± .740 | 2.68 ± .577 | $F(2,100) = 7.95$, $p < .001$ |
| Race/Ethnicity N | | | | |
| White | 12 | 32 | 31 | - |
| Black | 2 | 6 | 6 | - |
| Hispanic | 2 | 1 | 3 | - |
| Asian/Pacific Islander | 4 | 1 | 2 | - |
| More than 1 Race | 0 | 0 | 1 | - |

**Table S2. Correlations between D2R availability and summary-level experience sampling measures.** Note: no correlations remained statistically-significant after inclusion of participant covariates (age, sex, study sample, and difference in days between PET scan and EMA protocol start.

| | Ventral Striatum $BP_{ND}$ | Midbrain $BP_{ND}$ | Amygdala $BP_{ND}$ |
|---|---|---|---|
| Total # of Desires Reported | r = -0.067, p = 0.51 | r = -0.138, p = 0.163 | r = -0.086, p = 0.389 |
| Total # of Desire Resistance Attempts | r = 0.062, p = 0.534 | r = 0.161, p = 0.105 | r = 0.078, p = 0.437 |
| Total # of Desires Enacted | r = -0.094, p = 0.343 | r = -0.264, p = 0.007** | r = -0.102, p = 0.306 |
| Proportion of Attempts to Resist Made | r = 0.159, p = 0.108 | r = 0.273, p = 0.005** | r = 0.174, p = 0.079 |
| Proportion of Desires Enacted | r = 0.137, p = 0.169 | r = -0.071, p = 0.477 | r = 0.074, p = 0.457 |
| Mean Desire Rating | r = 0.040, p = 0.688 | r = 0.053, p = 0.592 | r = 0.035, p = 0.724 |
| Mean Goal Rating | r = 0.048, p = 0.239 | r = 0.203, p = 0.039* | r = 0.150, p = 0.130 |
| *** p < 0.001; ** p < 0.01; * p < 0.05. | | | |

**Table S3. Resistance attempt model comparison.** Regression coefficients are standardized. 95% confidence intervals appear below coefficients in brackets. Goal = degree of personal goal conflict, Desire = desire strength.

| | Intercept Only | Goal Only | Desire Only | Goal - Desire | Goal:Desire Ratio | Goal, Desire |
|---|---|---|---|---|---|---|
| Intercept | -0.69 ** | -0.95 *** | -0.83 *** | -0.96 *** | -0.94 *** | -0.96 *** |
| | [-1.14, -0.25] | [-1.41, -0.50] | [-1.21, -0.44] | [-1.41, -0.52] | [-1.40, -0.48] | [-1.43, -0.50] |
| Age | -0.55 *** | -0.60 *** | -0.52 *** | -0.59 *** | -0.61 *** | -0.61 *** |
| | [-0.81, -0.29] | [-0.90, -0.30] | [-0.77, -0.27] | [-0.88, -0.30] | [-0.92, -0.31] | [-0.91, -0.30] |
| Male | 0.33 | 0.43 | 0.35 | 0.42 | 0.45 | 0.44 |
| | [-0.09, 0.74] | [-0.07, 0.93] | [-0.08, 0.77] | [-0.07, 0.90] | [-0.06, 0.96] | [-0.07, 0.95] |
| Sample 1 | -0.81 | -0.62 | -0.46 | -0.58 | -0.62 | -0.64 |
| | [-1.74, 0.12] | [-1.38, 0.15] | [-1.10, 0.18] | [-1.33, 0.16] | [-1.40, 0.16] | [-1.42, 0.14] |
| Sample 2 | -0.55 * | -0.48 | -0.42 | -0.49 | -0.53 | -0.49 |
| | [-1.09, -0.01] | [-1.04, 0.08] | [-0.89, 0.06] | [-1.03, 0.06] | [-1.10, 0.05] | [-1.06, 0.08] |
| Goal | | 1.28 *** | | | | 1.29 *** |
| | | [1.20, 1.36] | | | | [1.21, 1.37] |
| Desire | | | -0.26 *** | | | -0.31 *** |
| | | | [-0.32, -0.20] | | | [-0.38, -0.23] |
| Goal-Desire | | | | 1.12 *** | | |
| | | | | [1.05, 1.20] | | |
| Goal:Desire | | | | | 1.27 *** | |
| | | | | | [1.19, 1.35] | |
| N (Survey) | 5752 | 5752 | 5752 | 5752 | 5752 | 5752 |
| N (Participant) | 103 | 103 | 103 | 103 | 103 | 103 |
| AIC | 6436.44 | 4981.82 | 6368.62 | 5380.32 | 5126.61 | 4915.15 |
| BIC | 6483.04 | 5028.42 | 6415.22 | 5426.92 | 5173.21 | 4968.41 |
| R2 (fixed) | 0.07 | 0.30 | 0.08 | 0.26 | 0.30 | 0.32 |
| R2 (total) | 0.29 | 0.52 | 0.30 | 0.48 | 0.52 | 0.54 |

*** $p < 0.001$; ** $p < 0.01$; * $p < 0.05$.

**Table S4. Resistance success model comparison.** Regression coefficients are standardized. 95% confidence intervals appear below coefficients in brackets. Goal = degree of personal goal conflict, Desire = desire strength.

| | Intercept Only | Goal Only | Desire Only | Goal - Desire | Goal:Desire Ratio | Goal, Desire |
|---|---|---|---|---|---|---|
| Intercept | -1.01 *** | -2.58 *** | -2.65 *** | -2.57 *** | -2.55 *** | -2.59 *** |
| | [-1.35, -0.66] | [-2.98, -2.18] | [-3.04, -2.25] | [-2.97, -2.16] | [-2.96, -2.15] | [-3.00, -2.18] |
| Age | -0.28 * | 0.02 | 0.04 | 0.02 | 0.02 | 0.02 |
| | [-0.51, -0.06] | [-0.23, 0.28] | [-0.21, 0.29] | [-0.24, 0.27] | [-0.24, 0.27] | [-0.24, 0.28] |
| Sex | 0.34 | 0.21 | 0.18 | 0.21 | 0.22 | 0.21 |
| | [-0.04, 0.71] | [-0.21, 0.63] | [-0.24, 0.59] | [-0.22, 0.63] | [-0.20, 0.65] | [-0.21, 0.64] |
| Sample 1 | -0.12 | 0.21 | 0.23 | 0.21 | 0.20 | 0.21 |
| | [-0.69, 0.46] | [-0.43, 0.85] | [-0.40, 0.86] | [-0.44, 0.86] | [-0.45, 0.85] | [-0.44, 0.86] |
| Sample 2 | -0.36 | -0.25 | -0.23 | -0.25 | -0.27 | -0.25 |
| | [-0.78, 0.06] | [-0.72, 0.23] | [-0.70, 0.23] | [-0.73, 0.23] | [-0.74, 0.21] | [-0.73, 0.22] |
| Resistance Attempt | | 3.31 *** | 3.63 *** | 3.26 *** | 3.26 *** | 3.26 *** |
| | | [3.10, 3.52] | [3.43, 3.83] | [3.06, 3.47] | [3.05, 3.46] | [3.05, 3.48] |
| Goal | | 0.21 ** | | | | 0.21 ** |
| | | [0.06, 0.35] | | | | [0.06, 0.36] |
| Resistance Attempt x Goal | | 0.20 * | | | | 0.24 * |
| | | [0.02, 0.38] | | | | [0.05, 0.42] |
| Desire | | | -0.32 *** | | | -0.32 *** |
| | | | [-0.43, -0.20] | | | [-0.44, -0.21] |
| Resistance Attempt x Desire | | | -0.03 | | | -0.07 |

| | | | | | | |
|---|---|---|---|---|---|---|
| | | | [-0.19, 0.12] | | | [-0.24, 0.09] |
| Conflict – Desire | | | | 0.40 *** | | |
| | | | | [0.27, 0.53] | | |
| Resistance Attempt x Goal – Desire | | | | 0.21 * | | |
| | | | | [0.03, 0.38] | | |
| Goal:Desire Ratio | | | | | 0.34 *** | |
| | | | | | [0.20, 0.47] | |
| Resistance Attempt x Goal:Desire Ratio | | | | | 0.18 * | |
| | | | | | [0.01, 0.35] | |
| N(Survey) | 5752 | 5752 | 5752 | 5752 | 5752 | 5752 |
| N (Participant) | 103 | 103 | 103 | 103 | 103 | 103 |
| AIC | 6485.01 | 4314.09 | 4313.41 | 4236.17 | 4269.69 | 4238.03 |
| BIC | 6524.96 | 4374.01 | 4373.33 | 4296.08 | 4329.60 | 4311.26 |
| R2 (fixed) | 0.03 | 0.43 | 0.44 | 0.44 | 0.44 | 0.44 |
| R2 (total) | 0.23 | 0.56 | 0.56 | 0.57 | 0.57 | 0.57 |

*** $p < 0.001$; ** $p < 0.01$; * $p < 0.05$.

**Table S5. Effects of social context on resistance attempt.** Regression coefficients are standardized. 95% confidence intervals appear beside coefficients in brackets. Goal = degree of personal goal conflict, Desire = desire strength.

| | Others Present Enacting Model |
|---|---|
| Intercept | -0.55 * [-1.04, -0.07] |
| Age | -0.61 *** [-0.92, -0.29] |
| Male | 0.43 [-0.10, 0.96] |
| Sample 1 | -0.64 [-1.45, 0.17] |
| Sample 2 | -0.50 [-1.10, 0.09] |
| Goal | 1.26 *** [1.15, 1.36] |
| Desire | -0.27 *** [-0.36, -0.18] |
| Others Present Enacting | -1.00 *** [-1.16, -0.84] |
| Goal x Others Present Enacting | -0.01 [-0.17, 0.14] |
| Desire x Others Present Enacting | -0.08 [-0.24, 0.07] |
| N | 5751 |
| N (SubjID) | 103 |
| AIC | 4769.60 |
| BIC | 4842.83 |
| R2 (fixed) | 0.35 |
| R2 (total) | 0.57 |

*** p < 0.001; ** p < 0.01; * p < 0.05.

**Table S6. Effects of social context on resistance success.** Regression coefficients are standardized. 95% confidence intervals appear below coefficients in brackets. Goal = degree of personal goal conflict, Desire = desire strength.

| | Others Present Enacting Model |
|---|---|
| Intercept | -2.37 *** [-2.84, -1.91] |
| Age | 0.11 [-0.14, 0.36] |
| Male | 0.23 [-0.17, 0.64] |
| Sample 1 | 1.13 * [0.22, 2.04] |
| Sample 2 | 0.09 [-0.44, 0.62] |
| Personal Goal | 0.11 [-0.07, 0.28] |
| Desire Strength | -0.35 *** [-0.48, -0.21] |
| Others Present Enacting | -1.27 *** [-1.56, -0.99] |
| Attempted to Resist | 3.07 *** [2.82, 3.32] |
| Attempted to Resist x Personal Goal | 0.27 * [0.05, 0.50] |
| Attempted to Resist x Desire Strength | -0.12 [-0.32, 0.08] |
| Attempted to Resist x Personal Goal x Others Present Enacting | -0.11 [-0.50, 0.29] |
| Attempted to Resist x Desire Strength x Others Present Enacting | -0.02 [-0.38, 0.33] |
| N | 5751 |
| N (SubjID) | 103 |
| AIC | 4080.22 |
| BIC | 4200.05 |
| R2 (fixed) | 0.49 |
| R2 (total) | 0.60 |

*** $p < 0.001$; ** $p < 0.01$; * $p < 0.05$.

**Table S7. Effects of dopamine D2 receptor availability on resistance attempt.** Regression coefficients are standardized. 95% confidence intervals appear beside coefficients in brackets. Goal = degree of personal goal conflict, Desire = desire strength.

| | VS | Midbrain | Amygdala |
|---|---|---|---|
| Intercept | -0.73 ** [-1.27, -0.20] | -0.82 ** [-1.38, -0.27] | -0.73 ** [-1.28, -0.19] |
| Age | -0.55 *** [-0.87, -0.24] | -0.61 ** [-0.98, -0.24] | -0.54 ** [-0.88, -0.19] |
| Male | 0.43 [-0.06, 0.93] | 0.42 [-0.09, 0.93] | 0.40 [-0.10, 0.90] |
| Sample 1 | -0.99 [-2.11, 0.12] | -1.09 [-2.23, 0.06] | -1.03 [-2.16, 0.10] |
| Sample 2 | -0.90 ** [-1.58, -0.22] | -0.64 [-1.33, 0.06] | -0.87 * [-1.57, -0.17] |
| PET-EMA Date Δ | -0.23 [-0.59, 0.14] | -0.21 [-0.58, 0.16] | -0.19 [-0.56, 0.18] |
| Goal | 1.30 *** [1.22, 1.39] | 1.29 *** [1.21, 1.37] | 1.29 *** [1.21, 1.37] |
| Desire | -0.30 *** [-0.38, -0.23] | -0.30 *** [-0.38, -0.23] | -0.30 *** [-0.38, -0.23] |
| D2R | 0.30 * [0.02, 0.57] | 0.07 [-0.25, 0.39] | 0.21 [-0.08, 0.50] |
| D2R x Goal | 0.15 *** [0.07, 0.23] | 0.02 [-0.06, 0.10] | 0.04 [-0.05, 0.12] |
| D2R x Desire | -0.06 [-0.13, 0.01] | -0.02 [-0.09, 0.06] | -0.05 [-0.12, 0.03] |
| N (Survey) | 5752 | 5752 | 5752 |
| N (Participant) | 103 | 103 | 103 |
| AIC | 4901.06 | 4921.35 | 4917.48 |
| BIC | 4980.95 | 5001.24 | 4997.37 |
| R2 (fixed) | 0.33 | 0.32 | 0.32 |
| R2 (total) | 0.54 | 0.53 | 0.54 |

*** p < 0.001;  ** p < 0.01;  * p < 0.05.

**Table S8. Effects of dopamine D2 receptor availability on resistance success.** Regression coefficients are standardized. 95% confidence intervals appear beside coefficients in brackets. Goal = degree of personal goal conflict, Desire = desire strength.

| | Ventral Striatum | Midbrain | Amygdala |
|---|---|---|---|
| Intercept | -2.90 *** [-3.37, -2.43] | -2.96 *** [-3.43, -2.50] | -2.88 *** [-3.35, -2.42] |
| Age | 0.14 [-0.12, 0.41] | 0.24 [-0.06, 0.54] | 0.17 [-0.12, 0.45] |
| Male | 0.25 [-0.16, 0.66] | 0.24 [-0.17, 0.64] | 0.24 [-0.17, 0.65] |
| Sample 1 | 1.15 * [0.23, 2.07] | 1.17 * [0.25, 2.09] | 1.15 * [0.23, 2.08] |
| Sample 2 | 0.07 [-0.50, 0.64] | 0.21 [-0.35, 0.77] | 0.06 [-0.51, 0.64] |
| PET-EMA Date Δ | 0.41 ** [0.11, 0.71] | 0.40 ** [0.10, 0.70] | 0.41 ** [0.11, 0.71] |
| Goal | 0.21 ** [0.06, 0.36] | 0.21 ** [0.06, 0.35] | 0.21 ** [0.07, 0.36] |
| Desire | -0.32 *** [-0.44, -0.21] | -0.33 *** [-0.45, -0.22] | -0.32 *** [-0.43, -0.20] |
| Resistance Attempt | 3.29 *** [3.07, 3.50] | 3.31 *** [3.10, 3.53] | 3.28 *** [3.06, 3.49] |
| Resistance Attempt x Goal | 0.23 * [0.05, 0.42] | 0.23 * [0.05, 0.42] | 0.24 ** [0.06, 0.42] |
| Resistance Attempt x Desire | -0.08 [-0.24, 0.08] | -0.07 [-0.23, 0.09] | -0.08 [-0.24, 0.08] |
| D2R | 0.16 [-0.10, 0.41] | 0.34 * [0.05, 0.62] | 0.19 [-0.08, 0.45] |
| D2R x Goal | -0.04 [-0.19, 0.12] | 0.04 [-0.10, 0.18] | 0.11 [-0.05, 0.27] |
| D2R x Desire | 0.03 [-0.09, 0.15] | 0.09 [-0.03, 0.20] | 0.07 [-0.06, 0.19] |
| D2R x Resistance Attempt | -0.17 [-0.37, 0.03] | -0.29 ** [-0.49, -0.08] | -0.15 [-0.37, 0.06] |
| D2R x Goal x Resistance Attempt | 0.07 [-0.12, 0.26] | -0.01 [-0.18, 0.16] | -0.19 [-0.38, 0.00] |
| D2R x Desire x Resistance Attempt | -0.02 [-0.18, 0.14] | -0.05 [-0.21, 0.11] | -0.06 [-0.24, 0.11] |
| N (Survey) | 5752 | 5752 | 5752 |
| N (Participant) | 103 | 103 | 103 |
| AIC | 4240.86 | 4233.55 | 4237.74 |
| BIC | 4360.69 | 4353.38 | 4357.57 |
| R2 (fixed) | 0.45 | 0.46 | 0.46 |
| R2 (total) | 0.57 | 0.57 | 0.57 |

*** p < 0.001; ** p < 0.01; * p < 0.05.

**Table S9. Effects of social context and dopamine D2 receptor availability on resistance attempt.** Regression coefficients are standardized. 95% confidence intervals appear beside coefficients in brackets. Goal = degree of personal goal conflict, Desire = desire strength.

| | VS | Midbrain | Amygdala |
|---|---|---|---|
| Intercept | -0.30 [-0.86, 0.27] | -0.39 [-0.97, 0.20] | -0.29 [-0.87, 0.28] |
| Age | -0.56 *** [-0.90, -0.23] | -0.61 ** [-1.00, -0.23] | -0.55 ** [-0.92, -0.19] |
| Male | 0.42 [-0.09, 0.94] | 0.41 [-0.12, 0.94] | 0.39 [-0.13, 0.92] |
| Sample 1 | -1.09 [-2.26, 0.07] | -1.16 [-2.36, 0.04] | -1.12 [-2.31, 0.06] |
| Sample 2 | -0.94 ** [-1.65, -0.22] | -0.67 [-1.40, 0.05] | -0.91 * [-1.64, -0.17] |
| PET-EMA Date delta | -0.26 [-0.64, 0.12] | -0.25 [-0.63, 0.14] | -0.23 [-0.61, 0.16] |
| Goal | 1.29 *** [1.18, 1.39] | 1.26 *** [1.15, 1.36] | 1.25 *** [1.15, 1.35] |
| Desire | -0.28 *** [-0.37, -0.18] | -0.27 *** [-0.36, -0.18] | -0.27 *** [-0.36, -0.18] |
| Others Present Enacting | -1.00 *** [-1.16, -0.84] | -1.01 *** [-1.17, -0.84] | -1.01 *** [-1.17, -0.84] |
| Personal Goal x Others Present Enacting | -0.04 [-0.20, 0.11] | -0.01 [-0.17, 0.15] | -0.01 [-0.16, 0.15] |
| Desire x Others Present Enacting | -0.08 [-0.23, 0.08] | -0.07 [-0.23, 0.08] | -0.09 [-0.24, 0.07] |
| D2R | 0.30 * [0.01, 0.59] | 0.07 [-0.27, 0.41] | 0.17 [-0.14, 0.48] |
| D2R x Goal | 0.23 *** [0.13, 0.34] | 0.04 [-0.06, 0.14] | 0.09 [-0.02, 0.20] |
| D2R x Desire | -0.09 * [-0.19, -0.00] | 0.04 [-0.05, 0.14] | -0.07 [-0.17, 0.03] |
| D2R x Others Present Enacting | -0.06 [-0.21, 0.10] | 0.03 [-0.14, 0.19] | 0.08 [-0.09, 0.24] |
| Goal x Others Present Enacting x D2R | -0.20 * [-0.36, -0.04] | -0.07 [-0.23, 0.09] | -0.13 [-0.29, 0.03] |
| Desire x Others Present Enacting x D2R | 0.09 [-0.06, 0.23] | -0.12 [-0.28, 0.03] | 0.06 [-0.09, 0.22] |
| N | 5751 | 5751 | 5751 |
| N (SubjID) | 103 | 103 | 103 |
| AIC | 4755.57 | 4778.09 | 4775.03 |
| BIC | 4875.40 | 4897.92 | 4894.86 |
| R2 (fixed) | 0.37 | 0.35 | 0.36 |
| R2 (total) | 0.57 | 0.57 | 0.57 |

*** $p < 0.001$; ** $p < 0.01$; * $p < 0.05$.

**Table S10. Effects of social context and dopamine D2 receptor availability on resistance success.** Regression coefficients are standardized. 95% confidence intervals appear beside coefficients in brackets. Goal = degree of personal goal conflict, Desire = desire strength.

| | Ventral Striatum | Midbrain | Amygdala |
|---|---|---|---|
| Intercept | -2.36 *** [-2.81, -1.91] | -2.44 *** [-2.88, -1.99] | -2.35 *** [-2.80, -1.91] |
| Age | 0.15 [-0.10, 0.40] | 0.27 [-0.01, 0.55] | 0.17 [-0.10, 0.44] |
| Male | 0.21 [-0.18, 0.60] | 0.19 [-0.19, 0.57] | 0.20 [-0.19, 0.58] |
| Sample 1 | 1.18 ** [0.30, 2.06] | 1.25 ** [0.39, 2.11] | 1.19 ** [0.32, 2.06] |
| Sample 2 | 0.07 [-0.47, 0.62] | 0.23 [-0.29, 0.76] | 0.08 [-0.46, 0.62] |
| PET-EMA Date delta | 0.38 ** [0.09, 0.67] | 0.39 ** [0.11, 0.67] | 0.39 ** [0.11, 0.68] |
| Attempted to Resist | 3.38 *** [3.15, 3.61] | 3.39 *** [3.16, 3.62] | 3.37 *** [3.14, 3.60] |
| Others Present Enacting | -1.44 *** [-1.71, -1.17] | -1.38 *** [-1.65, -1.11] | -1.40 *** [-1.67, -1.13] |
| Attempted to Resist x Others Present Enacting | 0.51 ** [0.15, 0.87] | 0.50 ** [0.14, 0.85] | 0.49 ** [0.13, 0.84] |
| D2R | 0.08 [-0.17, 0.34] | 0.44 ** [0.17, 0.71] | 0.12 [-0.14, 0.39] |
| D2R x Attempt | -0.14 [-0.36, 0.07] | -0.30 ** [-0.52, -0.09] | -0.13 [-0.36, 0.10] |
| D2R x Others Present Enacting | 0.26 [-0.02, 0.55] | -0.33 * [-0.60, -0.06] | 0.03 [-0.26, 0.33] |
| D2R x Attempt x Others Present Enacting | -0.05 [-0.41, 0.31] | 0.19 [-0.16, 0.54] | -0.02 [-0.41, 0.36] |
| N | 5751 | 5751 | 5751 |
| N (SubjID) | 103 | 103 | 103 |
| AIC | 4202.05 | 4195.29 | 4210.23 |
| BIC | 4295.25 | 4288.49 | 4303.43 |
| R2 (fixed) | 0.49 | 0.48 | 0.48 |
| R2 (total) | 0.59 | 0.58 | 0.58 |

*** $p < 0.001$; ** $p < 0.01$; * $p < 0.05$.

**Table S11. Resistance attempt model comparison including outlier observations.** Regression coefficients are standardized. 95% confidence intervals appear below coefficients in brackets. Goal = degree of personal goal conflict, Desire = desire strength.

| | Intercept Only | Goal Only | Desire Only | Goal - Desire | Goal:Desire Ratio | Goal, Desire |
|---|---|---|---|---|---|---|
| Intercept | -0.67 ** | -0.94 *** | -0.80 *** | -0.96 *** | -0.94 *** | -0.96 *** |
| | [-1.09, -0.25] | [-1.41, -0.48] | [-1.16, -0.43] | [-1.41, -0.52] | [-1.41, -0.47] | [-1.43, -0.49] |
| Age | -0.54 *** | -0.64 *** | -0.51 *** | -0.62 *** | -0.66 *** | -0.65 *** |
| | [-0.78, -0.29] | [-0.94, -0.33] | [-0.75, -0.27] | [-0.91, -0.33] | [-0.97, -0.35] | [-0.96, -0.34] |
| Male | 0.33 | 0.45 | 0.34 | 0.41 | 0.43 | 0.45 |
| | [-0.07, 0.72] | [-0.06, 0.95] | [-0.06, 0.74] | [-0.08, 0.90] | [-0.08, 0.95] | [-0.06, 0.97] |
| Sample 1 | -0.80 | -0.62 | -0.47 | -0.58 | -0.61 | -0.64 |
| | [-1.69, 0.08] | [-1.40, 0.15] | [-1.08, 0.15] | [-1.33, 0.17] | [-1.39, 0.17] | [-1.43, 0.15] |
| Sample 2 | -0.51 | -0.48 | -0.37 | -0.44 | -0.47 | -0.48 |
| | [-1.02, 0.01] | [-1.04, 0.09] | [-0.82, 0.08] | [-0.99, 0.11] | [-1.05, 0.10] | [-1.06, 0.10] |
| Goal | | 1.29 *** | | | | 1.30 *** |
| | | [1.21, 1.37] | | | | [1.22, 1.38] |
| Desire | | | -0.24 *** | | | -0.31 *** |
| | | | [-0.31, -0.18] | | | [-0.38, -0.24] |
| Goal-Desire | | | | 1.13 *** | | |
| | | | | [1.05, 1.21] | | |
| Goal:Desire | | | | | 1.25 *** | |
| | | | | | [1.17, 1.34] | |
| N (Survey) | 5820 | 5820 | 5820 | 5820 | 5820 | 5820 |
| N (Participant) | 103 | 103 | 103 | 103 | 103 | 103 |
| AIC | 6576.15 | 5131.32 | 6513.63 | 5480.98 | 5269.67 | 5061.47 |
| BIC | 6622.83 | 5178.00 | 6560.31 | 5527.67 | 5316.35 | 5114.82 |
| R2 (fixed) | 0.06 | 0.30 | 0.08 | 0.26 | 0.30 | 0.32 |
| R2 (total) | 0.27 | 0.53 | 0.28 | 0.49 | 0.52 | 0.54 |

*** p < 0.001; ** p < 0.01; * p < 0.05.

**Table S12. Resistance success model comparison including outlier observations.** Regression coefficients are standardized. 95% confidence intervals appear below coefficients in brackets. Goal = degree of personal goal conflict, Desire = desire strength.

| | Intercept Only | Goal Only | Desire Only | Goal - Desire | Goal:Desire Ratio | Goal, Desire |
|---|---|---|---|---|---|---|
| Intercept | -0.96 *** | -2.57 *** | -2.63 *** | -2.55 *** | -2.54 *** | -2.58 *** |
| | [-1.29, -0.64] | [-2.96, -2.17] | [-3.01, -2.24] | [-2.95, -2.15] | [-2.94, -2.14] | [-2.98, -2.18] |
| Age | -0.26 * | 0.04 | 0.07 | 0.03 | 0.03 | 0.03 |
| | [-0.47, -0.05] | [-0.21, 0.29] | [-0.17, 0.31] | [-0.22, 0.28] | [-0.22, 0.28] | [-0.22, 0.29] |
| Male | 0.32 | 0.21 | 0.16 | 0.20 | 0.21 | 0.21 |
| | [-0.03, 0.68] | [-0.20, 0.62] | [-0.24, 0.56] | [-0.22, 0.62] | [-0.20, 0.63] | [-0.21, 0.63] |
| Sample 1 | -0.12 | 0.22 | 0.25 | 0.22 | 0.21 | 0.22 |
| | [-0.66, 0.42] | [-0.41, 0.85] | [-0.36, 0.86] | [-0.42, 0.86] | [-0.43, 0.85] | [-0.42, 0.86] |
| Sample 2 | -0.34 | -0.25 | -0.22 | -0.25 | -0.27 | -0.26 |
| | [-0.74, 0.06] | [-0.72, 0.21] | [-0.67, 0.23] | [-0.72, 0.22] | [-0.74, 0.20] | [-0.73, 0.22] |
| Resistance Attempt | | 3.32 *** | 3.66 *** | 3.28 *** | 3.28 *** | 3.28 *** |
| | | [3.11, 3.53] | [3.47, 3.86] | [3.08, 3.48] | [3.07, 3.48] | [3.07, 3.48] |
| Goal | | 0.20 ** | | | | 0.21 ** |
| | | [0.07, 0.33] | | | | [0.08, 0.35] |
| Resistance Attempt x Goal | | 0.28 ** | | | | 0.30 *** |
| | | [0.10, 0.45] | | | | [0.13, 0.48] |
| Desire | | | -0.32 *** | | | -0.33 *** |
| | | | [-0.43, -0.20] | | | [-0.44, -0.21] |
| Resistance Attempt x Desire | | | -0.02 | | | -0.07 |
| | | | [-0.18, 0.13] | | | [-0.23, 0.09] |

| | | | | | | |
|---|---|---|---|---|---|---|
| Goal – Desire | | | | 0.40 *** | | |
| | | | | [0.27, 0.52] | | |
| Resistance Attempt x Goal – Desire | | | | 0.25 ** | | |
| | | | | [0.08, 0.43] | | |
| Goal:Desire Ratio | | | | | 0.35 *** | |
| | | | | | [0.22, 0.47] | |
| Attempted to Resist x Goal:Desire Ratio | | | | | 0.22 * | |
| | | | | | [0.05, 0.39] | |
| N (Survey) | 5820 | 5820 | 5820 | 5820 | 5820 | 5820 |
| N (Participant) | 103 | 103 | 103 | 103 | 103 | 103 |
| AIC | 6633.49 | 4360.72 | 4380.26 | 4283.96 | 4315.93 | 4283.71 |
| BIC | 6673.51 | 4420.74 | 4440.28 | 4343.98 | 4375.95 | 4357.07 |
| R2 (fixed) | 0.03 | 0.44 | 0.44 | 0.45 | 0.44 | 0.45 |
| R2 (total) | 0.20 | 0.56 | 0.56 | 0.57 | 0.57 | 0.57 |

*** $p < 0.001$; ** $p < 0.01$; * $p < 0.05$.

**Table S13. Effects of dopamine D2 receptor availability on resistance attempts including outlier observations.** Regression coefficients are standardized. 95% confidence intervals appear beside coefficients in brackets. Goal = degree of personal goal conflict, Desire = desire strength.

| | VS | Midbrain | Amygdala |
|---|---|---|---|
| Intercept | -0.72 ** [-1.26, -0.18] | -0.81 ** [-1.37, -0.24] | -0.72 * [-1.27, -0.17] |
| Age | -0.59 *** [-0.91, -0.26] | -0.64 *** [-1.01, -0.27] | -0.56 ** [-0.91, -0.21] |
| Male | 0.45 [-0.05, 0.95] | 0.42 [-0.10, 0.93] | 0.40 [-0.11, 0.91] |
| Sample 1 | -0.99 [-2.12, 0.13] | -1.07 [-2.24, 0.09] | -1.01 [-2.15, 0.13] |
| Sample 2 | -0.90 * [-1.59, -0.21] | -0.62 [-1.33, 0.09] | -0.86 * [-1.57, -0.16] |
| PET-EMA Date Δ | -0.23 [-0.59, 0.14] | -0.20 [-0.58, 0.17] | -0.18 [-0.55, 0.19] |
| Goal | 1.32 *** [1.24, 1.40] | 1.30 *** [1.22, 1.38] | 1.30 *** [1.22, 1.38] |
| Desire | -0.31 *** [-0.38, -0.23] | -0.30 *** [-0.38, -0.23] | -0.30 *** [-0.37, -0.23] |
| D2R | 0.30 * [0.03, 0.58] | 0.06 [-0.26, 0.39] | 0.22 [-0.07, 0.51] |
| D2R x Goal | 0.17 *** [0.09, 0.25] | 0.09 * [0.00, 0.17] | 0.10 * [0.02, 0.18] |
| D2R x Desire | -0.05 [-0.12, 0.02] | -0.02 [-0.10, 0.05] | -0.06 [-0.14, 0.01] |
| N (Survey) | 5820 | 5820 | 5820 |
| N (Participant) | 103 | 103 | 103 |
| AIC | 5044.55 | 5063.68 | 5058.19 |
| BIC | 5124.58 | 5143.71 | 5138.22 |
| R2 (fixed) | 0.34 | 0.32 | 0.32 |
| R2 (total) | 0.54 | 0.54 | 0.54 |

*** p < 0.001; ** p < 0.01; * p < 0.05.

**Table S14. Effects of dopamine D2 receptor availability on resistance success including outlier observations.** Regression coefficients are standardized. 95% confidence intervals appear beside coefficients in brackets. Goal = degree of personal goal conflict, Desire = desire strength.

| | Ventral Striatum | Midbrain | Amygdala |
|---|---|---|---|
| Intercept | -2.90 *** [-3.36, -2.44] | -2.95 *** [-3.41, -2.48] | -2.88 *** [-3.34, -2.42] |
| Age | 0.15 [-0.11, 0.41] | 0.26 [-0.04, 0.55] | 0.17 [-0.11, 0.45] |
| Male | 0.24 [-0.16, 0.65] | 0.23 [-0.17, 0.63] | 0.25 [-0.16, 0.65] |
| Sample 1 | 1.14 * [0.23, 2.06] | 1.17 * [0.26, 2.08] | 1.15 * [0.23, 2.06] |
| Sample 2 | 0.07 [-0.50, 0.63] | 0.20 [-0.35, 0.75] | 0.05 [-0.51, 0.62] |
| PET-EMA Date Δ | 0.40 ** [0.10, 0.70] | 0.39 ** [0.10, 0.68] | 0.40 ** [0.10, 0.70] |
| Goal | 0.19 ** [0.05, 0.34] | 0.22 ** [0.08, 0.36] | 0.20 ** [0.06, 0.34] |
| Desire | -0.33 *** [-0.44, -0.22] | -0.33 *** [-0.45, -0.22] | -0.32 *** [-0.43, -0.21] |
| Resistance Attempt | 3.31 *** [3.10, 3.52] | 3.32 *** [3.11, 3.53] | 3.29 *** [3.08, 3.50] |
| Resistance Attempt x Goal | 0.32 *** [0.14, 0.50] | 0.29 ** [0.11, 0.47] | 0.32 *** [0.14, 0.50] |
| Resistance Attempt x Desire | -0.07 [-0.23, 0.09] | -0.06 [-0.23, 0.10] | -0.07 [-0.23, 0.09] |
| D2R | 0.10 [-0.14, 0.35] | 0.34 * [0.06, 0.62] | 0.12 [-0.15, 0.38] |
| D2R x Goal | -0.13 [-0.28, 0.01] | 0.03 [-0.11, 0.17] | -0.06 [-0.19, 0.07] |
| D2R x Desire | 0.03 [-0.09, 0.15] | 0.08 [-0.03, 0.20] | 0.06 [-0.06, 0.18] |
| D2R x Resistance Attempt | -0.12 [-0.31, 0.08] | -0.28 ** [-0.48, -0.08] | -0.09 [-0.30, 0.12] |
| D2R x Goal x Resistance Attempt | 0.16 [-0.02, 0.35] | -0.01 [-0.18, 0.17] | 0.01 [-0.17, 0.18] |
| D2R x Desire x Resistance Attempt | -0.03 [-0.19, 0.13] | -0.06 [-0.21, 0.10] | -0.07 [-0.24, 0.11] |
| N (Survey) | 5820 | 5820 | 5820 |
| N (Participant) | 103 | 103 | 103 |
| AIC | 4284.45 | 4279.74 | 4286.69 |
| BIC | 4404.49 | 4399.79 | 4406.73 |
| R2 (fixed) | 0.46 | 0.47 | 0.46 |
| R2 (total) | 0.58 | 0.58 | 0.57 |

*** $p < 0.001$; ** $p < 0.01$; * $p < 0.05$.

**Table S15. Effects of social context on resistance attempt including outlier observations.** Regression coefficients are standardized. 95% confidence intervals appear beside coefficients in brackets. Goal = degree of personal goal conflict, Desire = desire strength.

| | Others Present Enacting |
|---|---|
| Intercept | -0.55 * [-1.05, -0.06] |
| Age | -0.65 *** [-0.97, -0.32] |
| Male | 0.45 [-0.09, 0.98] |
| Sample 1 | -0.64 [-1.46, 0.18] |
| Sample 2 | -0.50 [-1.10, 0.11] |
| Personal Goal | 1.26 *** [1.15, 1.36] |
| Desire Strength | -0.27 *** [-0.36, -0.18] |
| Others Present Enacting | -1.00 *** [-1.16, -0.84] |
| Goal x Others Present Enacting | -0.01 [-0.17, 0.15] |
| Desire x Others Present Enacting | -0.10 [-0.25, 0.05] |
| N | 5819 |
| N (SubjID) | 103 |
| AIC | 4910.59 |
| BIC | 4983.95 |
| R2 (fixed) | 0.34 |
| R2 (total) | 0.57 |

*** $p < 0.001$; ** $p < 0.01$; * $p < 0.05$.

**Table S16. Effects of social context on resistance success including outlier observations.** Regression coefficients are standardized. 95% confidence intervals appear beside coefficients in brackets. Goal = degree of personal goal conflict, Desire = desire strength.

| | Others Present Enacting |
|---|---|
| Intercept | -2.34 *** [-2.80, -1.88] |
| Age | 0.12 [-0.13, 0.37] |
| Male | 0.23 [-0.17, 0.63] |
| Sample 1 | 1.12 * [0.22, 2.02] |
| Sample 2 | 0.08 [-0.44, 0.61] |
| Personal Goal | 0.16 [-0.00, 0.33] |
| Desire Strength | -0.35 *** [-0.48, -0.21] |
| Others Present Enacting | -1.34 *** [-1.63, -1.06] |
| Attempted to Resist | 3.05 *** [2.81, 3.29] |
| Attempted to Resist x Personal Goal | 0.30 ** [0.08, 0.52] |
| Attempted to Resist x Desire Strength | -0.12 [-0.32, 0.08] |
| Attempted to Resist x Personal Goal x Others Present Enacting | 0.03 [-0.35, 0.40] |
| Attempted to Resist x Desire Strength x Others Present Enacting | -0.02 [-0.38, 0.33] |
| N | 5819 |
| N (SubjID) | 103 |
| AIC | 4124.21 |
| BIC | 4244.25 |
| R2 (fixed) | 0.50 |
| R2 (total) | 0.60 |

*** p < 0.001; ** p < 0.01; * p < 0.05.

**Table S17. Effects of social context and dopamine D2 receptor availability on resistance attempt including outlier observations.** Regression coefficients are standardized. 95% confidence intervals appear beside coefficients in brackets. Goal = degree of personal goal conflict, Desire = desire strength.

| | VS | Midbrain | Amygdala |
|---|---|---|---|
| Intercept | -0.28 [-0.85, 0.29] | -0.38 [-0.97, 0.21] | -0.28 [-0.86, 0.30] |
| Age | -0.60 *** [-0.93, -0.26] | -0.65 ** [-1.03, -0.26] | -0.57 ** [-0.94, -0.20] |
| Male | 0.45 [-0.08, 0.97] | 0.41 [-0.13, 0.94] | 0.39 [-0.14, 0.92] |
| Sample 1 | -1.10 [-2.28, 0.08] | -1.15 [-2.36, 0.07] | -1.10 [-2.29, 0.10] |
| Sample 2 | -0.95 ** [-1.67, -0.23] | -0.66 [-1.39, 0.08] | -0.90 * [-1.64, -0.16] |
| PET-EMA Date Δ | -0.27 [-0.65, 0.12] | -0.24 [-0.63, 0.16] | -0.21 [-0.60, 0.17] |
| Goal | 1.31 *** [1.20, 1.42] | 1.26 *** [1.15, 1.36] | 1.26 *** [1.16, 1.37] |
| Desire | -0.27 *** [-0.36, -0.18] | -0.26 *** [-0.35, -0.17] | -0.27 *** [-0.36, -0.17] |
| Others Present Enacting | -1.01 *** [-1.17, -0.85] | -1.00 *** [-1.16, -0.84] | -1.01 *** [-1.17, -0.84] |
| Goal x Others Present Enacting | -0.06 [-0.22, 0.10] | -0.01 [-0.17, 0.15] | -0.01 [-0.17, 0.15] |
| Desire x Others Present Enacting | -0.09 [-0.24, 0.06] | -0.09 [-0.24, 0.06] | -0.10 [-0.25, 0.06] |
| D2R | 0.31 * [0.02, 0.61] | 0.06 [-0.28, 0.41] | 0.18 [-0.13, 0.50] |
| D2R x Goal | 0.29 *** [0.18, 0.39] | 0.10 [-0.01, 0.21] | 0.18 *** [0.08, 0.28] |
| D2R x Desire | -0.08 [-0.17, 0.01] | 0.02 [-0.07, 0.11] | -0.09 [-0.18, 0.01] |
| D2R x Others Present Enacting | -0.08 [-0.24, 0.07] | 0.03 [-0.14, 0.19] | 0.06 [-0.10, 0.23] |
| Goal x Others Present Enacting x D2R | -0.30 *** [-0.47, -0.14] | -0.06 [-0.21, 0.10] | -0.21 * [-0.37, -0.04] |
| Desire x Others Present Enacting x D2R | 0.08 [-0.07, 0.22] | -0.09 [-0.24, 0.07] | 0.08 [-0.08, 0.24] |
| N | 5819 | 5819 | 5819 |
| N (SubjID) | 103 | 103 | 103 |
| AIC | 4886.25 | 4917.50 | 4906.77 |
| BIC | 5006.29 | 5037.54 | 5026.81 |
| R2 (fixed) | 0.37 | 0.35 | 0.35 |
| R2 (total) | 0.58 | 0.57 | 0.57 |

*** $p < 0.001$; ** $p < 0.01$; * $p < 0.05$.

**Table S18. Effects of social context and dopamine D2 receptor availability on resistance success including outlier observations.** Regression coefficients are standardized. 95% confidence intervals appear beside coefficients in brackets. Goal = degree of personal goal conflict, Desire = desire strength.

| | Ventral Striatum | Midbrain | Amygdala |
|---|---|---|---|
| Intercept | -2.34 *** [-2.77, -1.90] | -2.41 *** [-2.85, -1.98] | -2.33 *** [-2.77, -1.90] |
| Age | 0.18 [-0.07, 0.42] | 0.30 * [0.03, 0.57] | 0.19 [-0.07, 0.45] |
| Male | 0.19 [-0.18, 0.57] | 0.17 [-0.19, 0.54] | 0.18 [-0.19, 0.56] |
| Sample 1 | 1.19 ** [0.34, 2.04] | 1.26 ** [0.43, 2.10] | 1.20 ** [0.35, 2.04] |
| Sample 2 | 0.09 [-0.43, 0.61] | 0.24 [-0.26, 0.75] | 0.10 [-0.42, 0.62] |
| PET-EMA Date Δ | 0.38 ** [0.10, 0.66] | 0.39 ** [0.12, 0.66] | 0.39 ** [0.12, 0.67] |
| Attempted to Resist | 3.40 *** [3.18, 3.63] | 3.41 *** [3.18, 3.63] | 3.39 *** [3.16, 3.62] |
| Others Present Enacting | -1.45 *** [-1.73, -1.18] | -1.40 *** [-1.67, -1.13] | -1.41 *** [-1.68, -1.15] |
| Attempted to Resist x Others Present Enacting | 0.54 ** [0.19, 0.90] | 0.54 ** [0.19, 0.89] | 0.52 ** [0.17, 0.87] |
| D2R | 0.06 [-0.19, 0.31] | 0.44 ** [0.18, 0.71] | 0.08 [-0.18, 0.33] |
| D2R x Attempt | -0.12 [-0.33, 0.10] | -0.30 ** [-0.51, -0.08] | -0.09 [-0.31, 0.14] |
| D2R x Others Present Enacting | 0.28 * [0.00, 0.57] | -0.30 * [-0.57, -0.04] | 0.08 [-0.20, 0.37] |
| VS D2R x Attempt x Others Present Enacting | -0.08 [-0.44, 0.28] | 0.13 [-0.22, 0.47] | -0.07 [-0.44, 0.31] |
| N | 5819 | 5819 | 5819 |
| N (SubjID) | 103 | 103 | 103 |
| AIC | 4263.95 | 4255.53 | 4272.42 |
| BIC | 4357.31 | 4348.89 | 4365.79 |
| R2 (fixed) | 0.49 | 0.49 | 0.49 |
| R2 (total) | 0.59 | 0.58 | 0.58 |

*** $p < 0.001$; ** $p < 0.01$; * $p < 0.05$.

**Fig. S1. Effect of midbrain dopamine D2 receptor availability and resistance attempt on successful desire resistance.** The relationship between resistance attempt and successful resistance was dependent on individual differences in midbrain D2R availability ($\beta = -0.286$, CI [-0.489, -0.082], $Z = -2.76$, $p = 0.006$). Shaded areas represent ± 1 standard error.

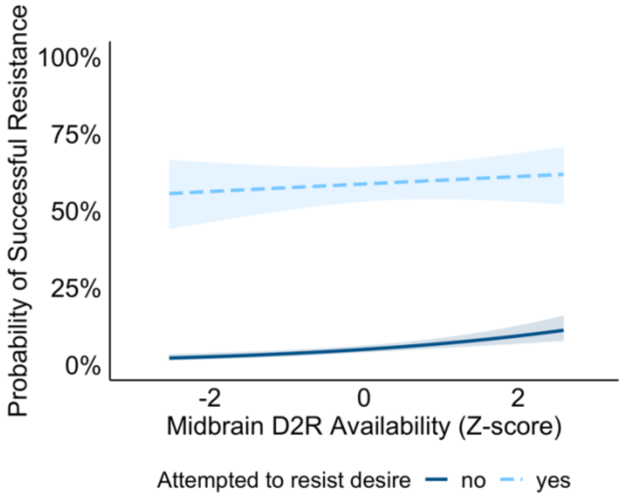
